# A subcellular atlas of *Toxoplasma* reveals the functional context of the proteome

**DOI:** 10.1101/2020.04.23.057125

**Authors:** Konstantin Barylyuk, Ludek Koreny, Huiling Ke, Simon Butterworth, Oliver M. Crook, Imen Lassadi, Vipul Gupta, Eelco Tromer, Tobias Mourier, Tim J. Stevens, Lisa M. Breckels, Arnab Pain, Kathryn S. Lilley, Ross F. Waller

## Abstract

Apicomplexan parasites cause major human disease and food insecurity. They owe their considerable success to novel, highly specialized cell compartments and structures. These adaptations drive their recognition and non-destructive penetration of host’s cells and the elaborate reengineering of these cells to promote growth, dissemination, and the countering of host defenses. The evolution of unique apicomplexan cellular compartments is concomitant with vast proteomic novelty that defines these new cell organizations and their functions. Consequently, half of apicomplexan proteins are unique and uncharacterized, and these cells are, therefore, very poorly understood. Here, we determine the steady-state subcellular location of thousands of proteins simultaneously within the globally prevalent apicomplexan parasite *Toxoplasma gondii*. This provides unprecedented comprehensive molecular definition to these cells and their novel compartments, and these data reveal the spatial organizations of protein expression and function, adaptation to hosts, and the underlying evolutionary trajectories of these pathogens.

## INTRODUCTION

Apicomplexa is a phylum of highly adapted unicellular eukaryotes specialized for intracellular parasitism in animals (Votýpka et al., 2017). The remarkable success of this group is evident by more than 6,000 described species that infect potentially every vertebrate and most invertebrates (Votýpka et al., 2017). Many apicomplexans cause devastating diseases in humans and livestock. Malaria, caused by *Plasmodium* spp., results in over 400,000 deaths and 200 million infections annually, with 3.2 billion people at risk (World Health Organization, 2018). Cryptosporidiosis (*Cryptosporidium* spp.) is the second leading cause of fatal infant diarrhea affecting 800,000 annually (Kotloff et al., 2013; Striepen, 2013). Toxoplasmosis (*Toxoplasma gondii*) occurs as chronic infections in ~30% of the human population and can cause life-threatening congenital toxoplasmosis, fetal malformation and abortion, blindness, and encephalitis (Havelaar et al., 2015). Furthermore, the economic damage of disease in livestock caused by apicomplexans is estimated in billions of US dollars annually (Rashid et al., 2019). Together these pathogens have a major effect on global health and prosperity, disproportionately affecting developing world regions.

Apicomplexans are deeply divergent organisms from better-studied model eukaryotic cell systems and, as parasites and pathogens, have displayed superb ingenuity for generation and specialization of new cell structures and compartments. For example, a dedicated apical structure enables non-destructive penetration and invasion of human and animal cells. This ‘apical complex’ includes a battery of different novel secretory compartments (micronemes, rhoptries, dense granules) for staged release of molecules required to search for, identify, penetrate and exploit the host’s cells (Figure 1*A*) (Kats et al., 2008; Lebrun et al., 2014). Apicomplexans have also developed novel gliding motility structures anchored in a pellicular cytoskeleton (Frénal et al., 2017). Furthermore, modified versions of two canonical endosymbiotic compartments, the mitochondrion and a remnant of a photosynthetic plastid (apicoplast), have developed in response to the metabolic needs of obligate parasitism (Sheiner et al., 2013).

**Figure 1.**
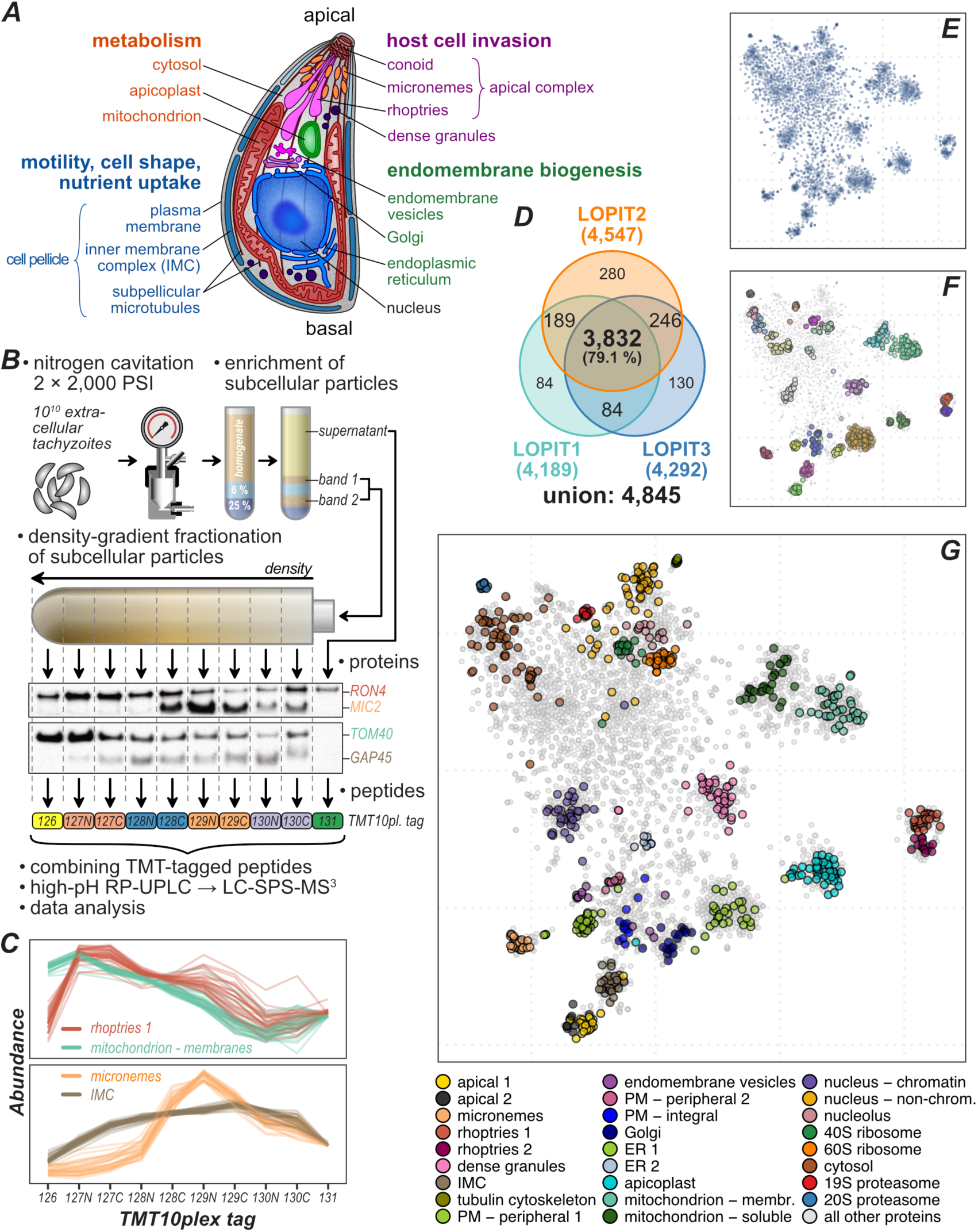
hyperLOPIT reveals organelle protein ensembles through measuring co-fractionation profiles of proteins. ***A.*** Schematic of *T. gondii* tachyzoite showing the main subcellular compartments and structures and their major functional roles. ***B.*** Summary of hyperLOPIT workflow applied to *T. gondii* extracellular tachyzoites. Cells are mechanically disrupted, homogenate fractionated (conditions optimization by Western blot, e.g. markers for rhoptries (RON4), micronemes (MIC2), mitochondria (TOM40) and IMC (GAP45)), and peptides labelled with a unique 10plex tandem mass tags for relative peptide quantitation by tandem mass spectrometry (LC-SPS-MS^3^). ***C.*** Abundance distribution profiles of select subcellular marker proteins measured in LOPIT2 experiment. Note the similarity with the WB results shown in panel ***B***. See **Figure S1** for concatenated profiles of all experiments (30plex). ***D.*** A Venn diagram showing the numbers of unique and shared proteins identified and quantified in all 10 fractions of the three hyperLOPIT experiments. ***E.*** A 2D-projection of the 30plex quantitative proteomic data (i.e., abundance distribution profiles) on 3,832 *T. gondii* proteins shared across three hyperLOPIT datasets. t-distributed stochastic neighbor embedding (t-SNE) was used for dimensionality reduction. Each data point represents an individual protein, and clustering of proteins reflects similarity of their abundance distribution profiles. ***F.*** Protein clusters discovered by the analysis of raw abundance distribution profiles with HDBSCAN overlaid on the t-SNE projection. Distinct clusters identified are indicated by color, and HDBSCAN clusters correspond to cluster centers identified by t-SNE. ***G.*** Mapping of 718 subcellular marker proteins on the t-SNE projection of *T. gondii* spatial proteome data.

Apicomplexans have not only created novel organization and biochemistry of their own cell compartments. Via secretion of complex mixtures of parasite proteins, they become active centers for subverting and remodeling the composition, organization, and properties of the host cells. Upon invasion of these host cells, they form, and typically remain within, a ‘parasitophorous vacuole’ decorated with secreted parasite proteins (Cesbron-Delauw et al., 2008). Parasite-secreted proteins also target and modify existing host cell compartments, including the nucleus, mitochondrion, ER, cytoskeleton, and plasma membrane. In doing so, they interfere with host control of defense and metabolism, cause reorganization of the host organelle positions and associations, change the mechanical properties of the host cell, and alter how infected cells interact with other human cells and tissues (Hakimi et al., 2017; Pernas et al., 2014; Soni et al., 2016). This exquisite redefinition of host cells reflects 100s of millions of years of co-evolution with their hosts and is orchestrated by the parasite secreted effector molecules delivered from the unique invasion machinery and compartments of these parasites. Moreover, this adaptation is ongoing with contemporary changes and variability that confounds adaptive immune responses and efforts to develop effective vaccines.

The novelty and divergence of apicomplexan cell compartments is concomitant with tremendous novelty of genes and proteins. This severely limits inferences of the cell biology of these organisms that can be made from knowledge of better-studied model organisms. Indeed, approximately half of apicomplexan proteins are known only as ‘hypotheticals’ and are unique to these cells (Swapna and Parkinson, 2017). Moreover, each apicomplexan lineage possesses its own new proteins, the products of ongoing adaptation to specific hosts (Woo et al., 2015). In such highly specialized cells, the subcellular location and function of proteins are tightly linked. Yet, despite decades of effort to understand the distribution of parasite proteins, typically relying on protein visualization by immunofluorescence microscopy (Woodcroft et al., 2012), the complete proteomes of most parasite compartments remain very poorly known. Even the locations of proteins of predicted function based on conserved sequences in other organisms are largely untested in apicomplexans. Thus, our understanding of the biological complexity and functional organization of these pathogens is severely limited by the lack of this basic information.

The roles and importance of apicomplexans’ unique protein repertoires are increasingly being screened for and tested using new tools that create resources such as genome-wide databases of mutant phenotypes, and single-cell expression profiles (Bushell et al., 2017; Sidik et al., 2016, 2018; Waldman et al., 2020). The lack of cellular context of much of the parasites’ proteomes, however, is a major bottleneck to interpreting these data. Here we address this critical deficiency, and the wider need to understand the compositional architecture of these deadly pathogens and its dynamics, by applying the spatial proteomic method hyperLOPIT (Christoforou et al., 2016; Mulvey et al., 2017) to capture the steady-state location of thousands of proteins in the apicomplexan *T. gondii*. This has provided an unprecedented, comprehensive understanding of the proteomic organization of an apicomplexan cell. This major advancement will have tremendous impact on apicomplexan research from basic cell biological processes to the discovery of novel therapeutic targets and strategies. Moreover, these data reveal the landscapes of cellular organization, function and evolution including gene expression programs, adaptative arms races with hosts, and the deeper evolutionary trajectories to parasitism, providing a new level of insight into these parasites.

## RESULTS

### Whole-cell biochemical fractionation of *Toxoplasma gondii* extracellular tachyzoites

To determine if the steady-state subcellular locations of thousands of proteins could be simultaneously captured in apicomplexans, we adapted the hyperLOPIT method for whole-cell spatial proteomics to *Toxoplasma gondii* tachyzoites, the extracellular form of this parasite that is primed for host cell invasion. The hyperLOPIT method exploits distinct abundance distribution profiles that organelles and subcellular structures form upon biochemical fractionation such as density-gradient centrifugation. Proteins exhibiting similar distribution profiles of abundance through these fractions are assigned to distinct subcellular structures (Christoforou et al., 2016; Mulvey et al., 2017).

To capture measurable variance of proteome profiles for hyperLOPIT assignment, optimization of both cell disruption methods and density gradient profiles were required. Using several subcellular marker proteins, western blot analysis of cell homogenates separated by differential and density-gradient centrifugation was used to determine suitable conditions (Figure 1*B*). Apicomplexan infectious zoites such as *Toxoplasma* tachyzoites have a robust cell pellicle (Figure 1*A*) that is resistant to simple cell disruption by hypotonic lysis. Nitrogen cavitation (Lacombe et al., 2019; Wang et al., 2014) was identified as the most effective, as well as non-heat-generating, method of cell disruption (**Table S1**). Membranous compartments and other cell particles were enriched from soluble cytosolic material by discontinuous density centrifugation of the homogenate, and this particulate material was then fractionated on continuous linear density gradients of iodixanol resulting in distinct enrichment profiles for a broad range of organelle markers (Figure 1*B*). The abundance distribution profiles of all detectable proteins were measured by sampling nine fractions across these gradients, plus one for the cytosol material fraction, labelling the peptides of each fraction with a unique TMT10plex isobaric tag, and quantifying relative peptide abundance across all fractions by mass spectrometry (Figures 1*C* and **S1**).

We performed three independent hyperLOPIT experiments, each with minor changes to cell rupturing, protein fraction preparation, and dispersal on density gradients, with the aim of maximizing captured resolvable differences amongst different subcellular protein niches (**Table S1**). In each experiment, we identified over 4,100 proteins with quantitative information across all 10 fractions (Figure 1*D*). 3,832 proteins were common to all three datasets providing complete abundance distribution profile information across 30 fractions (**Figure S1**, **Table S2**).

### HyperLOPIT assigns thousands of unknown proteins to subcellular niches

The protein fractionation data were analyzed for common abundance distribution patterns as evidence of protein association within subcellular niches (Breckels et al., 2016; Gatto et al., 2014). To visualize the 30-dimensional data, we used the machine-learning dimensionality reduction method t-distributed stochastic neighbor embedding (t-SNE) (van der Maaten et al., 2008). t-SNE projections indicated the presence of complex structure in the data with proteins resolved into multiple distinct clustered sets (Figure 1*E*). To verify that the clusters displayed in the t-SNE projection were not artefacts of modelling, we analyzed the untransformed data with the unsupervised clustering algorithm ‘hierarchical density-based spatial clustering of applications with noise’ (HDBSCAN) (Campello et al., 2013). The clusters found in the untransformed data by HDBSCAN corresponded to the cores of many of the clusters observed in the t-SNE map (Figure 1*F*). To assess if these protein clusters represent genuine biological protein assemblages, we compiled a set of 656 known marker proteins belonging to cell organelles compartments, structures or substructures based either on previous location studies, or strong evidence of protein function (**Table S3**). When projected onto t-SNE maps, these markers sort according to the clusters providing strong evidence that the hyperLOPIT-derived clusters do represent cell niches (Figure 1*G*). These clusters represent all known apicomplexan compartments or subcompartments, demonstrating that hyperLOPIT produced a highly resolved proteomic map of the *T. gondii* tachyzoite. The resolution of these data discerns membranous organelles (e.g. mitochondrial, micronemes, ER, Golgi), cytoskeletal elements (e.g. inner membrane complex, apical complex structures), molecular complexes (e.g. ribosome and proteasome subunits) and subcompartmental organization (e.g. outer and inner peripheral, and integral plasma membrane proteins).

To test the veracity of the hyperLOPIT clusters, 62 proteins associated with clusters representing distinct organelles or subcellular structures were epitope-tagged by endogenous gene fusion and the location of the protein determined by immuno-fluorescence microscopy. All 62 proteins were previously either completely uncharacterized, or in some cases had provisional annotation apparently in conflict with their hyperLOPIT-inferred location. All of the reporter-tagged proteins showed subcellular location consistent with their hyperLOPIT predictions, further supporting the very high correlation of the hyperLOPIT cluster data with subcellular niche (Figure 2 and **S2**).

**Figure 2.**
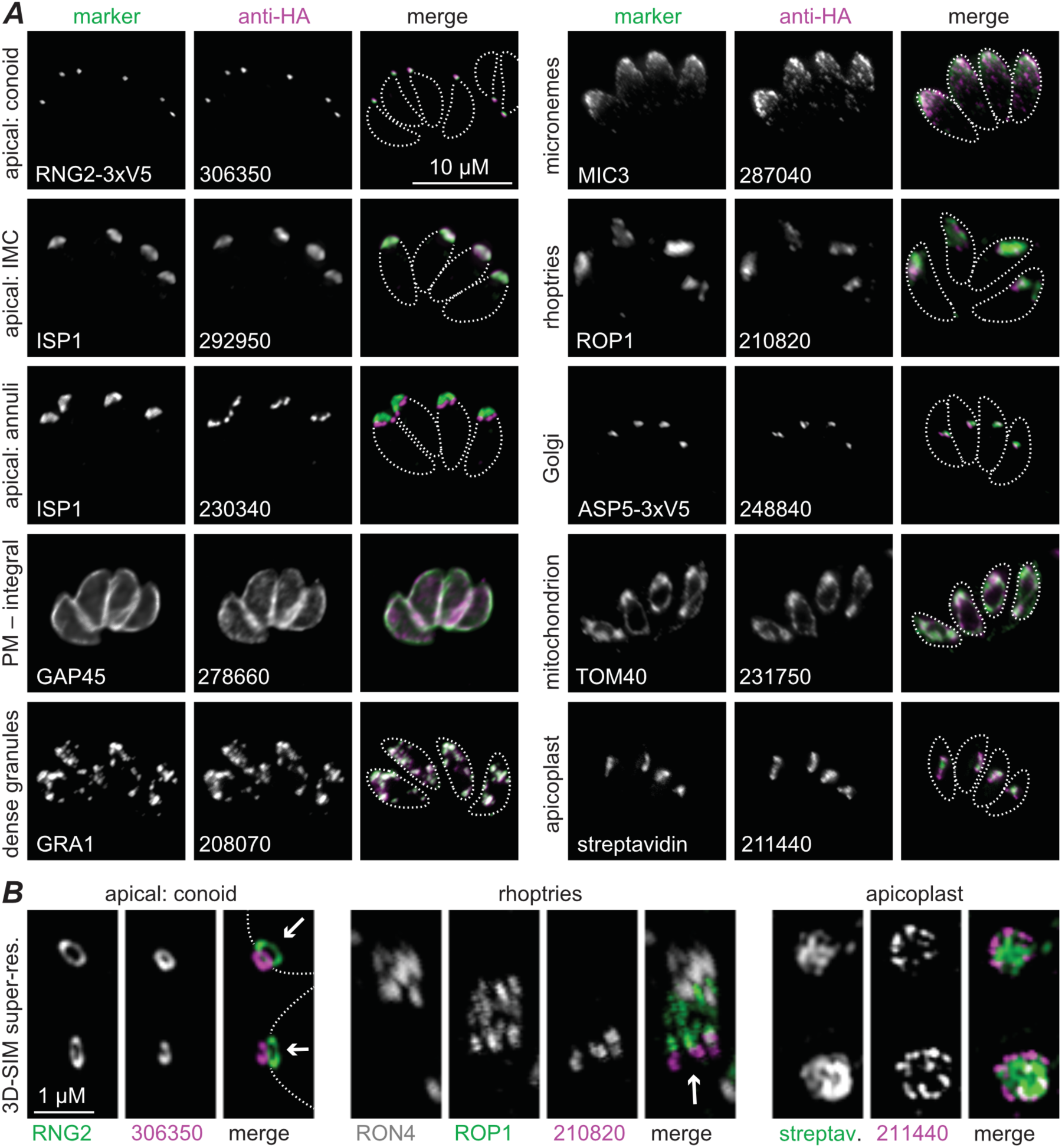
Validation of hyperLOPIT-predicted subcellular locations. ***A.*** Examples of uncharacterized proteins epitope-tagged and detected by immunofluorescence microscopy (magenta) co-located with named marker proteins (green). Cell outlines are indicated (dashed lines) in the merge. See **Figure S2** for all validated proteins. Scale bar = 10 µm for all. ***B.*** Optical super-resolution (3D-SIM) images of select proteins (magenta) from ***A*** co-located with subcellular marker proteins (green). Arrows indicate the cell posterior-to-anterior cell axis. Scale bar = 1 µm for all.

The resolution of protein clusters allows the prediction of subcellular location of all detected proteins by supervised machine-learning methods using the marker-protein distributions. The 62 newly validated proteins were added to the previous 656 markers to give 718 markers defining 26 distinct subcellular niches. We analyzed the data by a recently developed Bayesian classification method based on t-augmented Gaussian mixture models (TAGM) to probabilistically assign proteins to a set of defined classes (Crook et al., 2018, 2019). This method performs similarly to the established modelling methods (e.g. support vector machine, k-nearest neighbors, random forest) (Crook et al., 2018), but has the advantage of calculating a membership probability uniformly to all classes. This is achieved by estimates of the posterior probability of protein allocation to one of the defined subcellular classes or an outlier component which accounts for the noise in the data.

The expectation-maximization algorithm was used to compute *maximum a posteriori* (MAP) estimates of the TAGM model parameters from the known 718 marker proteins. Using these models, we analyzed the abundance distribution profiles of the remaining 3,114 proteins and obtained the probability of every protein belonging to the respective most likely subcellular class of the defined 26 and not being an outlier. We applied a uniform localization probability cut-off of 99% across all the 26 subcellular classes to retain only high-confidence protein classification results (Figures 3*A, B* and **S3**, **Table S4**). Of the 3,832 proteins measured across all three independent hyperLOPIT experiments, we assigned 1,916 proteins of previous unknown location (50% of the input) to one of 26 subcellular niches with a localization probability above 99%. The remaining 1,198 proteins, amounting to 31.3% of the detected proteins, are not assigned to any one location by TAGM-MAP approach with sufficient confidence (Figure 3*B*, *unassigned*).

**Figure 3.**
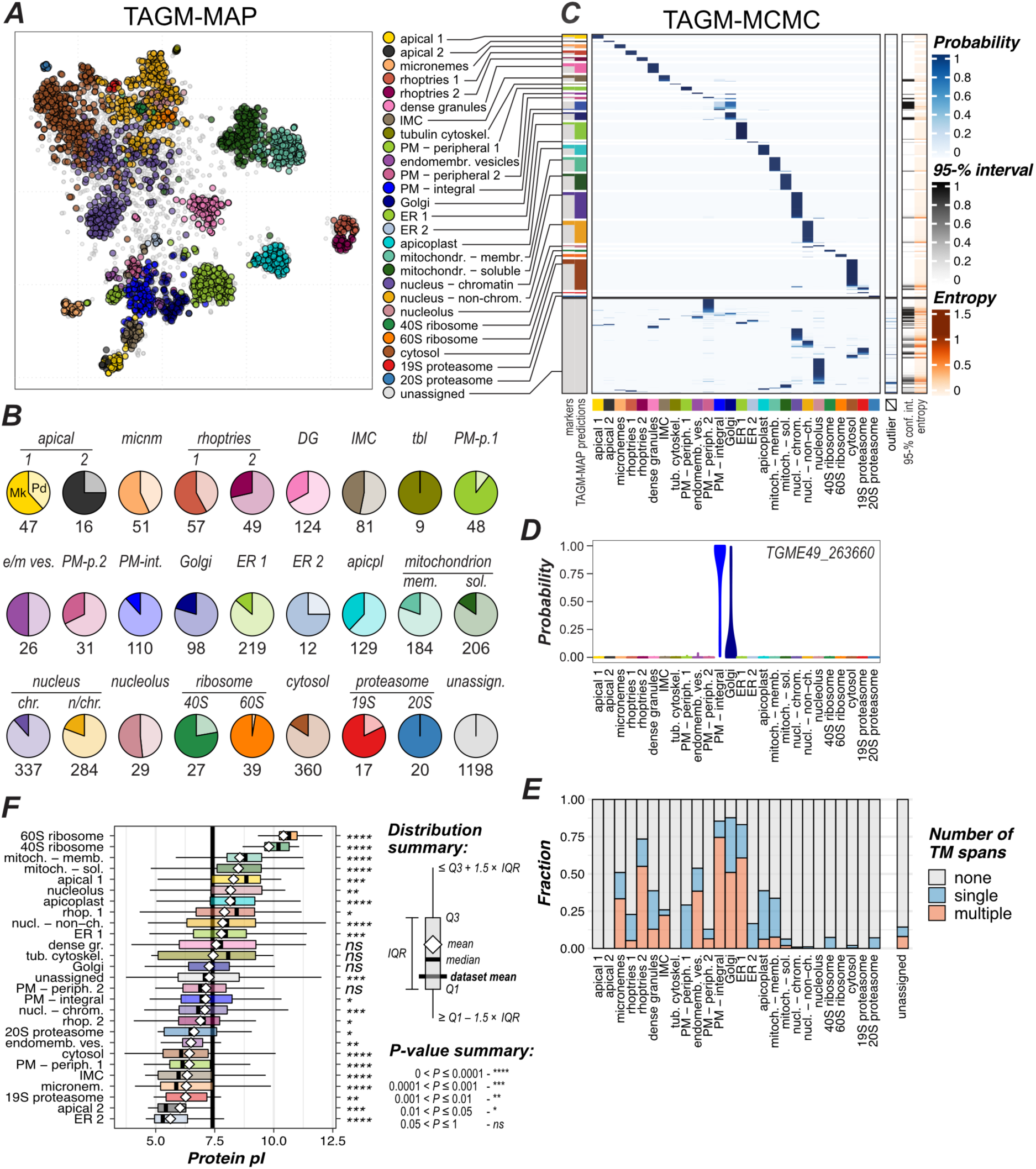
Protein assignment to known subcellular niches by a supervised Bayesian classification. ***A.*** TAGM-MAP predicted steady-stage location of proteins (99% probability) superimposed on the t-SNE projection of the 30plex hyperLOPIT data for 3,832 proteins. ***B.*** Number of proteins assigned to each location. Marker proteins (*Mk*: previously characterized proteins + verified proteins as in Figures 2, **S2**) are indicated in dark color, newly assigned protein predictions (*Pd*: at 99% TAGM-MAP probability) in light color. ***C.*** Heatmap showing proteins ordered by the TAGM-MAP-assigned class (rows) against joint probabilities of proteins to belong to each of the 26 defined subcellular classes or the outlier component (columns) inferred by TAGM-MCMC. Colorbars on the right show the uncertainty of TAGM-MCMC localization as the 95% equitailed confidence interval of the TAGM-MCMC localization probability (in shades of grey) and the mean Shannon entropy (in shades of red). ***D.*** A violin plot showing an example (TGME49_263660) TAGM-MCMC distribution of localization probabilities across the 26 subcellular niches. The most probable location predicted by TAGM-MAP and TAGM-MCMC for this protein is *PM – integral*, but there is also significant probability of localization to *Golgi*, consistent with signals seen for proteins that might cycle between multiple compartments. ***E.*** Fractions of monotopic and polytopic integral membrane proteins (blue and red, respectively) by subcellular class. ***F.*** Compartment-specific distributions of protein charge (computed pI) are shown as Tukey box plots (legend at right). The probability of class-specific means differing from the dataset average by chance is shown to the right. See also **Figures S3** and **S4**.

Steady-state determination of protein locations in a population of cells overlooks the dynamic behaviors that many proteins can have, including regulated location changes, trafficking intermediates, organelle contact points, and proteins with dual or multiple locations. Occupation of multiple locations by a protein will manifest as a composite abundance distribution profile in the hyperLOPIT data. To test if these dynamic protein behaviors can be detected, we sampled from the entire distribution of posterior location probabilities for each protein across all modelled subcellular niches using a fully Bayesian TAGM analysis employing Markov-chain Monte Carlo (MCMC) methods (Crook et al., 2018) (Figures 3*C, D* and **Table S5**). Most TAGM-MAP-assigned protein locations correspond to single, high-probability location associations by TAGM-MCMC, consistent with steady-state single locations for these proteins (Figure 3*C*). Some TAGM-MAP-assigned compartments, however, show enrichment of proteins with probability distributions across multiple compartments by TAGM-MCMC. For instance, many integral plasma membrane proteins show elevated probability for the Golgi, as do endomembrane vesicle proteins (Figure 3*C*, *D*). By contrast, the secretory organelles of the apical complex (micronemes, rhoptries, dense granules), which are also destination compartments of the secretory pathway, are dominated by single TAGM-MCMC assignments. This is consistent with a dynamic bidirectional exchange of proteins between Golgi, vesicles, and plasma membrane, whereas the proteomes of rhoptries, micronemes, and dense granules once established are static in this lifeform of *T. gondii*. Thus, TAGM-MCMC is apparently able to capture some of the dynamic properties of the *T. gondii* spatial proteome.

The TAGM-MCMC analysis also allows uncertainty quantification in the subcellular location of proteins; in particular, those that are *unassigned* proteins by TAGM-MAP (Figure 3*C*). In the TAGM-MAP model the majority of these proteins have a high probability of belonging to the outlier component (**Figure S3**), whereas TAGM-MCMC reports high probabilities of most of them belonging to a subcellular class. Many of these proteins are attributed to nuclear and cytosolic components, which could indicate their trafficking between these niches. However, we are cautious with this interpretation because of limitations of the subcellular fractionation method used in maintaining nuclear and cytosolic integrity. The remaining TAGM-MAP *unassigned* proteins are attributed to one of the defined subcellular classes with greater uncertainty (Figure 3*C*) which might indicate dynamic location behaviour of these proteins.

### HyperLOPIT achieves extensive apicomplexan cell compartment proteomic resolution

Interpretation of the cellular resolution of *T. gondii* achieved by hyperLOPIT requires deciphering the manner of the physical disruption and separation of organelles and subcellular structures. This, in turn, provides knowledge of proteins’ and compartments’ physical associations with one another and, thus, new insight into the biochemical organization of the cell.

Clear definition of distinct membrane-bound compartments (e.g. mitochondrion, apicoplast, rhoptries, micronemes, dense granules, ER) indicates that these structures were separated from one another relatively intact, and this has provided confident identification of respective proteomes. There is evidence also of some rupturing of mitochondria and rhoptries. Both organelles resolved as two clusters with an enrichment for integral membrane proteins in one and a depletion of membrane-anchored proteins in the other (Figures 3*A*, *E* and **S4*A***). This resolution is consistent with distinct abundance distribution profiles formed by each population of proteins: 1) the membrane-attached cohort dispersing with the organelle membranes only; and 2) the soluble cohort sharing a composite of the membrane profile of intact organelles and the distribution of released soluble proteins for ruptured organelles. This serendipitous distinction provided a further level of organelle proteome resolution and knowledge: proteins associated either directly or indirectly with membranous components of the organelle; and organelle soluble proteins and complexes.

The inner membrane complex (IMC) is a distinctive feature of apicomplexans that is comprised of a proteinaceous membrane skeleton that supports flattened membranous cisternae appressed to the cytosolic face of the plasma membrane (Figure 1*A*). The IMC is an essential platform for motility during invasion, maintaining cell shape and organization, and formation of new cells during cytokinesis (Harding and Meissner, 2014). A major IMC cluster resolved separately from plasma membrane clusters (Figures 3 and **S1**) indicating some level of dissociation of the IMC from the plasma membrane, and the proteins that associate with each, during cell rupturing. The membranous cisternae of the IMC are arranged in a tiled pattern that line the majority of the cell body including a single conical cisterna that occupies the apical portion (~10%) of the cell — the so-called apical cap (Figure 1*A*). Known apical cap proteins and proteins that locate to a series of small rings or ‘annuli’ at its posterior boundary resolved separately from IMC proteins seen in the rest of the cell (Figures 2, 3). This indicates dissociation at this boundary during cell disruption and a stronger attachment of the annuli structures to the apical cap than the posterior IMC cisternae. The apically resolved proteins also include all known proteins associated with the conoid and apical polar ring, two invasion-related structural components of the cell’s apical extremity (Figure 2 and **S2**). These apical proteins resolved further as two clusters, *apical 1* and *2*, although, this does not appear to represent a spatial differentiation, but rather a biochemical differentiation (Figures 3, **S1**). *Apical 1* is enriched in proteins with basic pI whereas *apical 2* with acidic pI (Figures 3*F* and **S4*C***), suggesting that biophysical interactions contribute to the hyperLOPIT resolution of these cells. Finally, the majority of the cell’s tubulin occurs in a basket of microtubules that underlie and support the IMC (Figure 1*A*). Tubulins, however, resolved with a select group of known subpellicular microtubule-associated proteins (MAPs) separately from either the IMC or apical proteins as a fourth cluster (Figure 3) indicating their dissociation from the proteinaceous subpellicular network of the IMC. Thus, the IMC as a definitive complex component of the apicomplexan cell pellicle resolved into four hyperLOPIT clusters of substructural associations.

The plasma membrane proteome resolved as three biochemically distinct clusters enriched in integral membrane proteins (*PM – integral*), peripheral proteins on the external leaflet dominated by the members of GPI-anchored surface antigen glycoprotein (SAG)-related sequence (SRS) protein family (Jung et al., 2004) (*PM – peripheral 1*), and peripheral proteins on the internal/cytosolic leaflet (*PM – peripheral 2*) (Figure 3, **Table S4**). ER proteins show subcompartment resolution also (Figure 3). A major class of ER proteins (*ER 1*) is enriched in integral membrane proteins. A second small group of more acidic, soluble proteins forms a distinct cluster (*ER 2*) that includes heat shock proteins (BiP, Hsp90, and DnaK family protein), and several other proteins implicated in protein folding and processing (Figures 3*E-F*). This provides novel insight into subcompartment organization in the ER of these parasites. The abundance distribution profiles of *ER 2* proteins are more similar to those of the *apicoplast* rather than *ER 1* (**Figure S1**), suggesting some degree of association between these two. Given that most apicoplast proteins traffic through the ER, this association might reflect a role of these proteins in folding and redox processes during sorting of proteins to the apicoplast (Biddau et al., 2018). Indeed, BiP was recently found among proteins pulled-down by an apicoplast-residing thioredoxin TgATrx2 (Biddau et al., 2018).

Our implementation of the hyperLOPIT method was tailored to fractionate and resolve subcellular membranous niches, notably associated with invasion and host interaction. However, cytosolic large protein complexes, such as the proteasome and ribosome subunits, stand out from the rest of cytosolic proteins and are, in fact, among the tightest and best resolved clusters (Figures 3 and **S1**). Evidence of additional structure in these regions of the hyperLOPIT maps (Figure 1*E-F*) indicates further resolution of protein associations in these complex spaces.

### Compartment proteomes provide massive expansion of knowledge of apicomplexan subcellular complexity

Of the 1,916 proteins that the hyperLOPIT could assign to known compartments with strong support, 795 (41.5%) were previously designated as ‘hypothetical proteins’, 335 (17.5%) annotated only as conserved domain- or repeat-containing proteins, 256 (13%) annotated as generic functions such as ‘transporter’ or ‘… family protein’, and for 228 (12%) their assigned function is ‘putative’ (Gajria et al., 2007). Hence, for only 16% (302) of these proteins was there any more certain notion of the protein’s functional annotation, typically assigned through protein similarity to conserved eukaryotic proteins, and the majority of these still lacked identified and/or experimentally validated locations. The hyperLOPIT assignments of protein location in *Toxoplasma*, therefore, provide an enormous advancement in our knowledge of protein composition of subcellular compartments and niches (Figure 3*B*).

Protein compartments that mediate parasite-host interaction is a facet of apicomplexan biology with tremendous expansion of knowledge provided by hyperLOPIT. Three distinct secretory compartments deliver proteins either onto the parasite surface, directly into the host cytoplasm, or into membranous compartments that the parasite occupies within its host cell. These proteins are delivered from micronemes, rhoptries, and dense granules, and they facilitate essential parasite processes: extracellular motility and host attachment (micronemes); penetration and invasion of the host cell (rhoptries); and manipulation of host defenses, metabolism, and acquisition of nutrients from this host (rhoptries and dense granules) (Lebrun et al., 2014). The importance of these functions to parasite infection, virulence and disease has focused much research attention on these compartments and their protein cargo. For each of micronemes, rhoptries and dense granules, 29, 47, and 41 proteins, respectively, had been previously identified. HyperLOPIT identifies a further 22, 59, and 83 proteins to each of these three compartments, providing extensive new knowledge of the proteins that mediate interactions with the host (Figure 3*B* and **Table S4**). Of these new proteins, 22, 43 and 49 from the three respective organelles lacked apparent signal peptides that might otherwise have predicted their location to secretory organelles. We tested 15 of these signal peptide-lacking proteins and verified that all locate to their assigned organelles (Figures 2 and **S2**, **Table S3**).

The separation of rhoptries into two distinct clusters provides a new understanding of the cell biological division within this organelle: *rhoptries 1* enriched with soluble cargo; *rhoptries 2* enriched with proteins associated with organelle maintenance and biogenesis, even capturing maturase processes (e.g. aspartyl protease 3) of the final steps of protein sorting to rhoptries (**Table S4a**) (Dogga et al., 2017). Furthermore, new insight is gained into the spatial organization of these organelles. Rhoptries have been known to partition select proteins into the anterior tapered rhoptry ‘neck’ from those in the posterior ‘bulb’, and this separation correlates with timing of secretion and function: neck proteins during host penetration, bulb proteins managing the subsequent infection. Our locating of new rhoptry proteins by microscopy revealed further subdomains within rhoptries, with some proteins exclusive to the posterior base of the bulb, and others marking both the anterior and posterior rhoptry extremities (Figures 2 and **S2**).

The parasite surface is also a critical site of interaction with the host. For the parasite, it is the site to transport material into and out of the cell and sense environmental changes that it must respond to. For the host, the parasite surface is a display of foreignness and invasion against which it can mount immune defense. The GPI-anchored SAG proteins are the best-known surface molecules (Jung et al., 2004), but relatively few integral membrane proteins in the plasma membrane that act as receptors, transporters or managing plasma membrane properties and functions are known. The cluster of integral plasma membrane proteins (*PM – integral*) contains 110 proteins, providing great expansion of knowledge of this proteome (Figure 3). Of nine proteins from this cluster that we gene-tagged with reporters, two were visible at the cell surface, but seven were below detectable limits by IFA suggesting that many of these proteins are of very low abundances. This is consistent with low abundance of receptors and transporters in other cell systems, and the challenge for their discovery by other protein location methods.

### hyperLOPIT resolves the cellular landscapes of proteome expression, function, adaption and evolution within the parasite cell

The differential behaviors and programs of apicomplexan organelles and structures that drive protein regulation, function, adaptation and evolution can only be resolved using comprehensive proteome samples of the different cell compartments. The hyperLOPIT spatial proteome of *Toxoplasma* provides the necessary statistical power to assess these cell properties for the first time in an apicomplexan.

#### Some, but not all, compartments show tight transcriptional regulatory control

In *Toxoplasma*, previous efforts to identify new candidate proteins for select compartments have used the correlation of transcript abundance profiles across the cell cycle making the assumption that co-located proteins are co-expressed (Bai et al., 2018; Lacombe et al., 2019; Long et al., 2017; Sheiner et al., 2011). However, an objective assessment of the coordination of gene expression for compartment proteins has not been previously possible without comprehensive knowledge of the spatial distribution of the proteome. To test for compartment-correlated transcriptional control, we collated a wide range of quantitative transcriptomic data and compared within-cluster correlations of co-expression to that between a cluster and the rest of the cell proteome (Figure 4*A*). In several clusters there is strong support for within cluster co-expression (Figures 4*B* and **S5**, **Table S6**). The genes for large protein complexes show particularly strong coordinated expression: apical complex, *19S* and *20S proteasome* subunits, *40S* and *60S ribosomes*. Membrane compartments for host invasion and interaction — micronemes, rhoptries, and dense granules — also show strong coordinated expression, as do the apicoplast and the IMC, although with less support. Other compartment-wide proteomes showed either smaller or no evidence of coordinated expression (e.g. soluble mitochondrion proteins).

**Figure 4.**
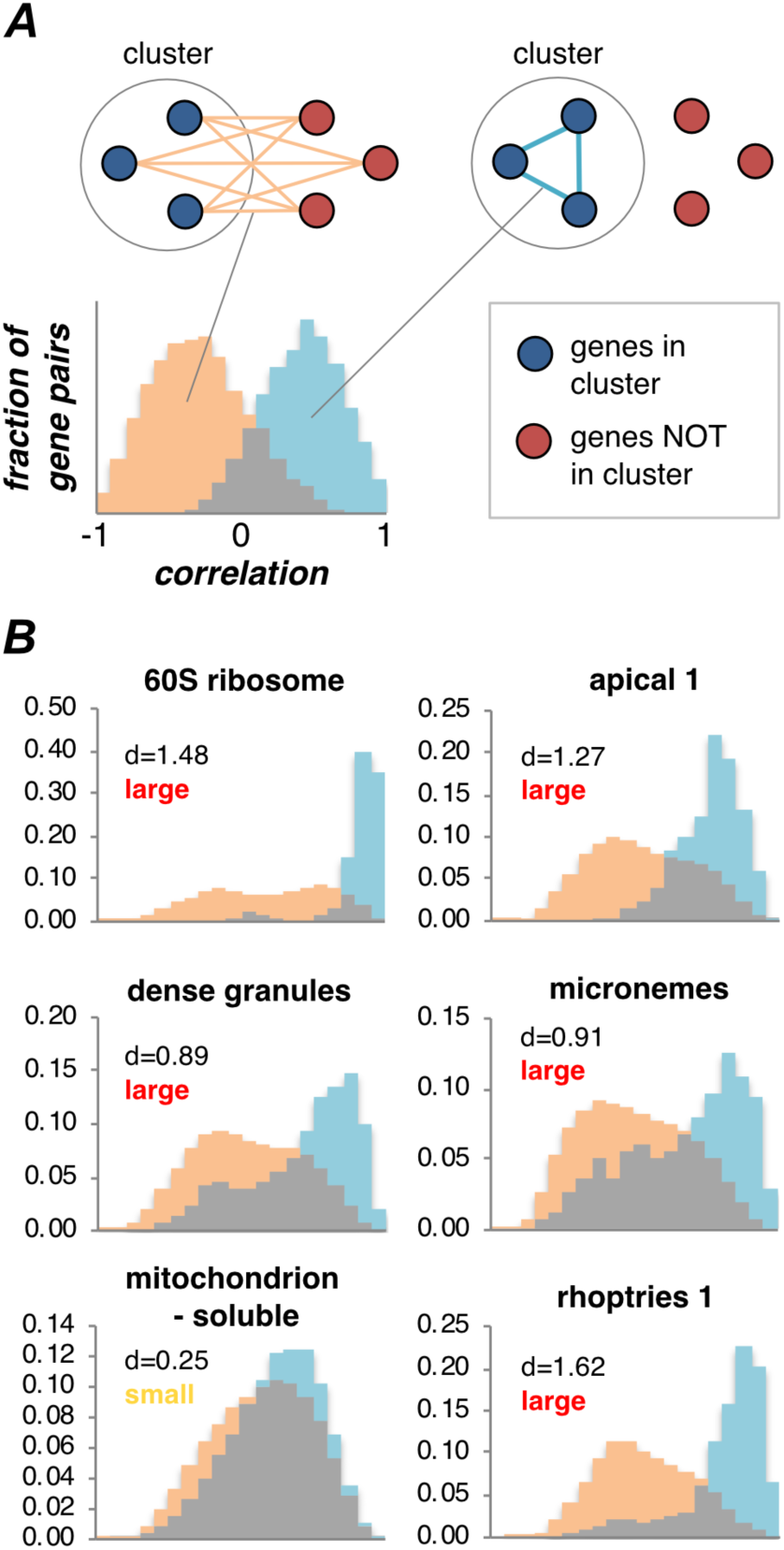
Correlation of gene expression patterns within subcellular compartments. ***A.*** Schematic of analysis of gene co-expression according to protein location. The distribution of co-expression levels between members of a cluster (blue) is plotted against this distribution between members of the cluster and all other genes (orange). ***B.*** Gene co-expression levels for select hyperLOPIT clusters measured as Pearson correlations. Cohen’s *d* values are shown above each chart along with *effect size* descriptors. See also **Figure S5** for all cluster results.

#### Subcellular proteomes reveal biophysical and functional partitioning of the cell

Protein properties are adapted to the environments and processes in which they operate. Proteomic data from subcellular niches, therefore, can report on the biochemical conditions of these microenvironments across cell compartments and programs. Protein pI values often reflects the pH of their local environment, and clear differences are seen in average pI values of proteins in the different subcellular niches (Figures 3*F* and **S4*C***, **Table S7a**). This includes the stepwise acidification through the secretory pathway: ER, Golgi to endomembrane vesicles. It also reveals an apparently basic pH of the apicoplast, a property of this organelle that was previously not known, and which could indicate a role of pH in protein import.

Our data also report differences in trans-membrane trafficking programs and membrane properties throughout the cell. The Sec61 complex is a common entry point for protein into the endomembrane system from which they are sorted to multiple destinations, many central to host interactions. Co-translational ER-import is mediated by interactions of the ER import machinery with cleavable N-terminal signal peptides. Comparison of signal peptides from *T. gondii* and their apicomplexan orthologues for different endomembrane niches reveal compositional differences between protein groups destined to different locations (Figures 5*A* and **S6**, **Table S8b**). These data imply a level of sorting selection even at these early stages of protein synthesis. Membrane-spanning proteins interact with the lipid bilayers that they are embedded into. Analysis of the distribution of lengths of apicomplexan single-span protein trans-membrane domains shows clear differences between compartments (Figure 5*B*). These differences most likely reflect lipid compositional difference across the cell that might also govern protein-sorting events. For instance, microneme proteins share similar long membrane spans with the plasma membrane consistent with this being the destination of microneme proteins once secreted. By contrast, dense granule proteins do not follow the increase in transmembrane domain seen from early to late parts of the secretory pathway (Sharpe et al., 2010). Dense granule proteins must avoid insertion into the parasite plasma membrane post secretion and their trend for shorter membrane spans might contribute to their onward trajectory into the host.

**Figure 5.**
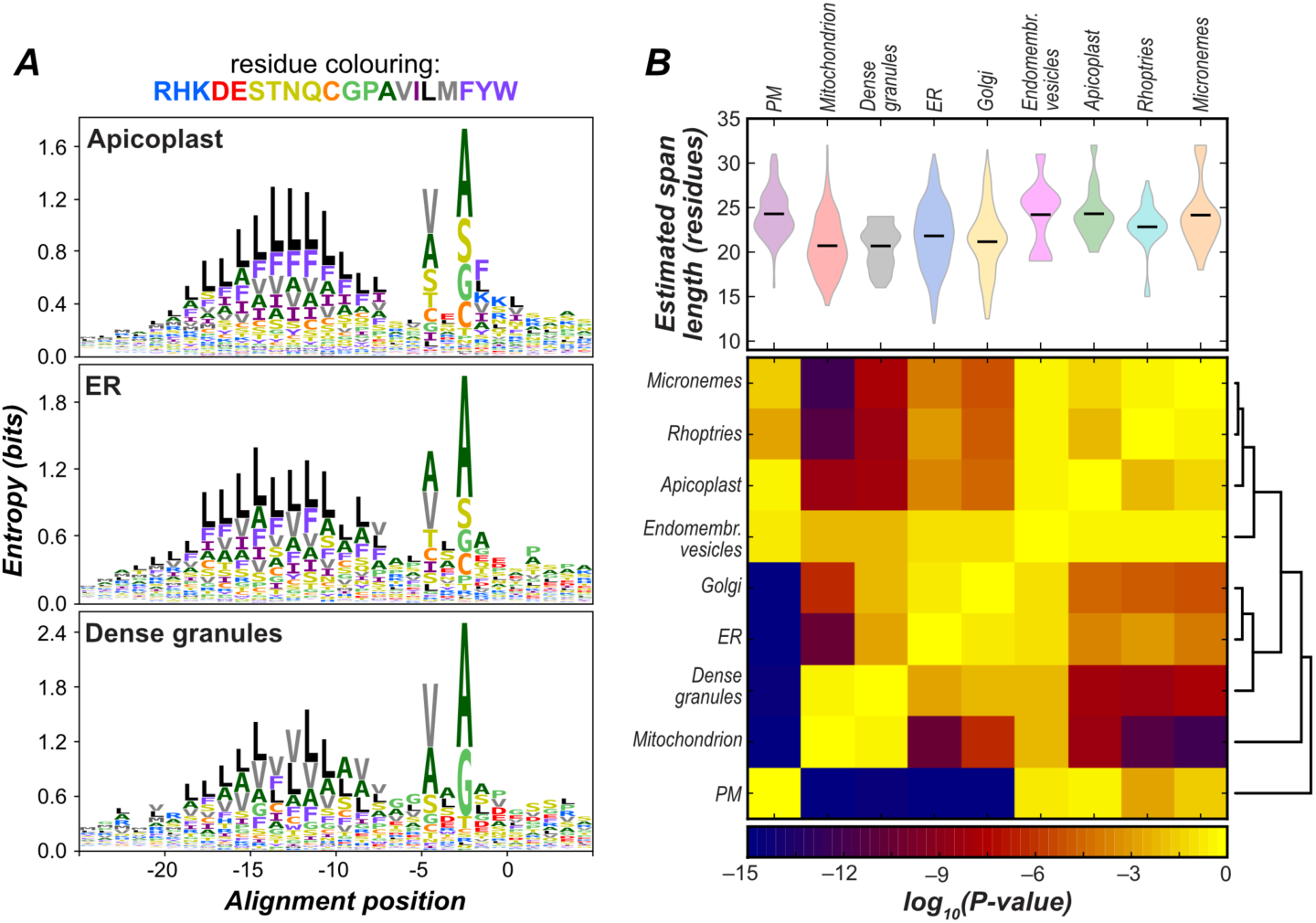
Distinction of properties of apicomplexan signal peptide (A) and transmembrane domain sequences (B) according to subcellular compartment. ***A.*** Differences in relative positional abundances of amino acids for signal peptide (SP) sequences of proteins from apicomplexan endomembrane compartments shown as logo plots anchored on the cleavage site (position 0). See also **Figure S6** and **Table S8a,b**. Amino acids colored by physicochemical properties. ***B.*** Distributions of apicomplexan transmembrane (TM) span length for single-span proteins of different compartment. The length distributions (violin plots) were compared pairwise by Mann-Whitney *U* test, and the resulting *p*-values (heatmap) were used to cluster membrane type. See also **Table S8c**.

The relative redundancy of proteomes across the subcellular landscape was also assessed using our extensive representation of compartment proteomes. Data from a genome-wide CRISPR-Cas9 knockout screen in *T. gondii* was employed where phenotype was measured during *in vitro* tachyzoite propagation (Sidik et al., 2016). Combining this genetic screen, with unambiguous evidence of protein expression in tachyzoites, enables the uneven compartment distribution of relatively dispensable versus indispensable proteins to be seen (Figures 6*A* and **S7*A***, **Table S7d**). The plasma membrane (including *PM-integral*), dense granules, micronemes, rhoptries, and the IMC show the largest bias for dispensable proteins in these conditions (Figure 6*A*), indicating a high incidence of expressed protein function redundancy. These compartments, therefore, are apparently unable to follow the otherwise common trend of parasite gene loss and complexity minimalization. By contrast, the apicoplast shows a paucity of dispensable proteins (Figure 6*A*). Thus, despite this organelle being a remnant of a former photosynthetic lifestyle, and its early interpretation as “evolutionary baggage”, it is now clear that this has become a highly reduced organelle supported by a bare essential proteome. Other examples of compartments with a high proportion of indispensable proteins include the mitochondrion, nucleus, proteasome, and ribosome (Figure 6*A*).

**Figure 6.**
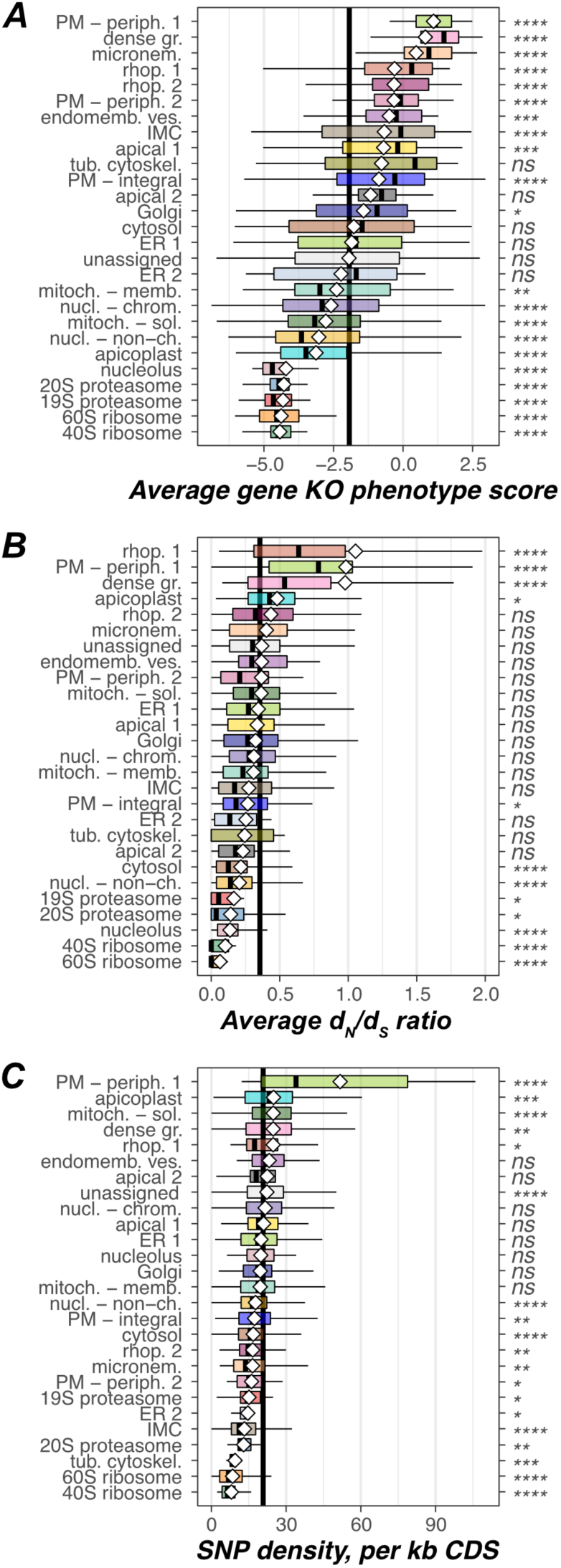
*T. gondii* subcellular compartments show distinct distributions of the functional redundancy of the proteomes (A), selection pressure (B), and genetic polymorphism (C). ***A.*** Compartment-specific distribution of protein functional redundancy expressed as the average gene knockout (KO) phenotype score quantifying the contribution of each *T. gondii* gene to the parasite fitness during in vitro culture (a negative score indicates relatively indispensable genes; a positive score indicates dispensable genes). ***B.*** Compartment-specific distributions of evolutionary selection pressures expressed as protein-average ratio of non-synonymous and synonymous mutation rates (*d*_N_/*d*_S_ ratio). ***C.*** Compartment-specific distributions of genetic polymorphism expressed as the density of single nucleotide polymorphisms (SNP) per kilobase of gene coding sequence (CDS). Compartment-specific distributions are shown as Tukey box plots as for Figure 3*F*. See also **Figure S7**.

#### Heterogeneous compartment host-adaptive responses

Parasites of humans and animals operate under enormous selective pressures to successfully target, exploit, replicate and then transmit within available hosts – all under constant surveillance and attack by the host immune system. As a zoonotic infectious agent, *T. gondii* is also adapted to exploiting a variety of different warm-blooded organisms. The strength and nature of selective pressures on a protein is evident by the ratio of rates of non-synonymous (*d*_*N*_) to synonymous (*d*_*S*_) point mutations for a gene, and the distribution of protein *d*_*N*_/*d*_*S*_ values inform on the within-cell distribution of these pressures and parasite responses across compartments.

Gene single nucleotide polymorphism (SNP) properties were analyzed across the subcellular compartments from population data for 62 *T. gondii* geographical isolates (Figures 6*B, C* and **S7*B, C***, **Tables S*7b, c***) (Lorenzi et al., 2016). Compartments with highly positive-skewed *d*_*N*_/*d*_*S*_ distributions are those of the external peripheral plasma membrane proteins, the soluble content of rhoptries, and the dense granules (Figure 6*B*). This implies strong positive selection for change, but also a high capacity of proteins in these niches to tolerate changes. Such proteins within these compartments are likely at the frontline of host-pathogen interaction and adaptation. In stark contrast to the high rate of change of the peripheral external plasma membrane proteins is that of the integral plasma membrane proteins which are biased for purifying selection (low *d*_*N*_/*d*_*S*_) (Figure 6*B*). These differences reveal the tension between exposure to host immune factors and maintenance of plasma membrane function, and likely contribute to driving the expression of these proteins down. Other cell niches under purifying selection are those for central cellular function: ribosomes, cytosol, non-chromatin nuclear proteins, nucleolus, and proteasome (Figure 6*B*).

SNP density within coding sequences also responds to compartment evolution, and for many compartments SNP density correlates with *d*_*N*_/*d*_*S*_ (e.g. both high for plasma membrane peripheral proteins, low for ribosomes) (Figure 6*C*). An unexpected mutation behavior, however, is observed with the mitochondrial soluble proteins that show significant enrichment for higher than average SNP densities (Figure 6*C*), but no increase in *d*_*N*_/*d*_*S*_ (Figure 6*B*). This enrichment for synonymous, or ‘silent’, mutations indicates selection for codon usage changes across strains. This likely has implications for translation efficiency differences and metabolic flux control in this important metabolic compartment. A similar bias for SNP density is seen in the apicoplast also, although here some selection for protein sequence change is also seen (Figure 6*B, C*). Thus, modulation of metabolic control might be an important driver of host tissue and/or taxon preference, or even virulence across parasite populations.

#### Compartment-specific evolutionary trajectories to parasitism in Apicomplexa

A resolved apicomplexan spatial proteome also allows the broader evolution of apicomplexan parasites to be assessed. We asked the question: when in the evolution of these parasites did different cell compartments and functions display the greatest rates of innovation? We surveyed the distribution of new protein orthologues across cell compartments over phylogenetic distance (Figure 7 and **Tables S9*a-f***). These data show that different cell compartments display very different rates of evolutionary protein innovation. At the most ancient level of the last eukaryotic common ancestor (LECA), as expected, orthologues are enriched across core cellular compartments, including the cytosol and complexes for protein expression, sorting and turnover (Figure 7). By contrast, the compartments most enriched for recent, coccidian-specific orthologues include the dense granules, rhoptry soluble fraction, micronemes, conoid and peripheral surface proteins — all components of the cell that define the interaction with its hosts. Dense granules show greatest novelty and are apparently most instrumental to the recent evolution in *Toxoplasma* and its close relatives. Other cellular locations show earlier, apicomplexan-specific accelerated evolution that have likely been important to the adaptation of apicomplexans as parasites: the IMC, which is key to parasite motility, host contact and invasion; and the ‘*nucleus – chromatin*’ cluster, which is consistent with evolution of novel gene regulatory networks shared by parasites (Woo et al., 2015). Chromerids are apicomplexans’ closest photosynthetic relatives and they also live in association with animal communities (Janouškovec et al., 2013). Innovation in the integral plasma membrane proteome, notably enriched in membrane transporters, is seen in these group’s common ancestors that might indicate the beginning of molecular exchange with animal partners. Even deeper, rapid change in the mitochondrion is evident prior to divergence of apicomplexans from dinoflagellate algae, and this is consistent with many known peculiarities of this essential metabolic organelle shared by both groups (Danne et al., 2013; Waller and Jackson, 2009). And finally, in the common ancestor of the Alveolata, the group that includes ciliates and is defined by the pellicle organization of sub-plasma membrane alveolae (IMC), enrichment for new inner leaflet peripheral plasma membrane proteins is seen. These proteins include several Ca^2+^ and cGMP receptive molecules (e.g. Calcium Dependent Protein Kinase 3, Protein Kinase G) of the signaling cascades that are central to apicomplexan invasion and host egress events. This is the first biochemical evidence for the common coupling of this cell ultrastructure with this critical function so early in apicomplexan evolution. Collectively, these data provide an unprecedented view of the evolutionary chronology of apicomplexan cells and their trajectory to parasitism.

**Figure 7.**
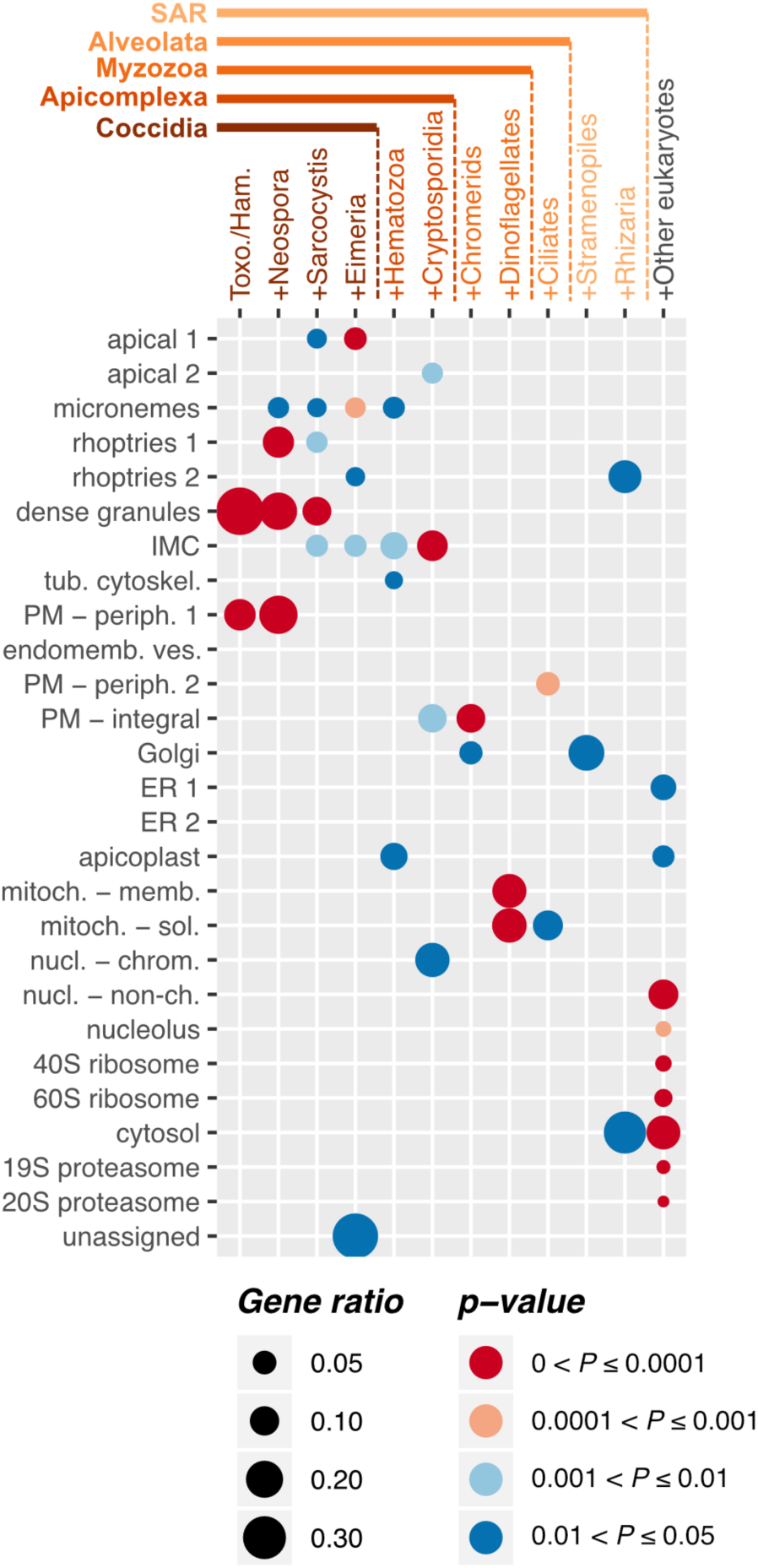
*T. gondii* subcellular compartment proteomes reveal the tempo of compartment evolution over evolutionary time. A dot plot showing the distribution of significant enrichments for new protein orthologues at twelve phylogenetic distance levels within hyperLOPIT-defined apicomplexan compartment classes. *p*-values (colors) calculated by under-representation hypergeometric test, and scaled according to the gene ratio (fraction of novel proteins in a compartment against all novel proteins at a given phylogenetic distance level). Toxo./Ham. = *Toxoplasma*/ *Hammondia*; SAR = Stramenopiles/Alveolata/Rhizaria. See also **Tables S9a-f**.

## DISCUSSION

The application of hyperLOPIT to the *T. gondii* extracellular tachyzoite provides the first comprehensive high-resolution spatial proteomic map of an apicomplexan cell. This is the first application of this method to a non-model eukaryotic providing sub-cellular niche composition of standard and novel compartments on an unprecedented level. Further, these data uncover a myriad of layers of new insight into the biochemical, functional and evolutionary organisation of these major human pathogens. Overall, we identified and quantified 3,832 proteins across three independent hyperLOPIT experiments and assigned 2,634 proteins to 26 distinct subcellular niches with 99% Bayesian posterior localization probability. These include virtually all known apicomplexan cell compartments including those specific to these parasites. The most immediate outcome of our study is a massive expansion of known organelle proteomes. In the case of invasion organelles – micronemes, rhoptries, and dense granules – this provides tremendous new knowledge of the complexity of effector repertoires secreted into the host upon parasite invasion, growth, and egress. We also capture the proteins involved in the biogenesis and maintenance of these organelles that otherwise have been overlooked due to the challenges of their discovery and bias towards studying their secreted contents. Major new elements of the proteomes of the apical subdomain of the IMC and the apical cytoskeletal structures including the conoid have also been discovered, both structures central to host invasion. Since these sub-cellular niches are phylogenetically restricted to apicomplexans, the empirical definition of their biochemistry is essential to understand their functions. Furthermore, with hundreds of proteins assigned to the mitochondrion and apicoplast, the metabolic capacity and activities of these otherwise enigmatic endosymbiotic organelles in parasites can now be addressed with far greater depth.

Because hyperLOPIT is independent from any inferences of protein function and/or location from conserved domains, sequence motif prediction or orthologues in other model organisms, it does not suffer from potential pitfalls of these approaches. For example, TGME49_310290 is annotated as a ‘regulator of chromosome condensation (RCC1) repeat-containing protein’ based on sequence similarity, suggesting a nuclear location for this protein. This protein, however, was assigned to the *mitochondrion – solubl*e cluster by hyperLOPIT and this location validated by microscopy (**Figure S2**). Similarly, the annotated ‘rhoptry kinase family proteins’, ROPs, suggest rhoptry localization, as is indeed the case for many such proteins (e.g., ROP11, 20, 24, 26) (Lebrun et al., 2014). However, several ‘ROP’ proteins (ROP32-35: TGME49_270920, TGME49_201130, TGME49_240090, and TGME49_304740) were attributed to dense granules by hyperLOPIT, and recently confirmed experimentally for ROP34 and ROP35 (Beraki et al., 2019). Moreover, proteins can relocate between compartments over evolutionary time, further confounding orthology-based inferences. For example, apicomplexans have expanded a family of mitochondria-targeted RNA-binding proteins (RAPs) with *Toxoplasma* encoding 16 such RAP-domain proteins (Lee and Hong, 2004; Woo et al., 2015). Most were assigned to the *mitochondrion – soluble* cluster by hyperLOPIT as expected. One RAP protein (TGME49_211890), however, that is a product of a Sarcocystidae-specific gene duplication, was assigned to the apicoplast and this location was also verified (**Figure S2**). While the organellar function of RAP proteins awaits discovery, this relocation of a mitochondrial protein to the apicoplast indicates evolutionary transfer of function between cell compartments. Such transfers might also account for ‘ROPs’ being dispersed in rhoptries and dense granules as parasite-host interactions continue to evolve.

There are a number of reasons why a fraction of proteins has not been confidently assigned to a subcellular compartment. Firstly, as a supervised machine learning algorithm, TAGM is unable to model unknown subcellular niches or those that are poorly understood for a lack of sufficient known proteins to serve as markers. Proteins that occur in such cellular niches will be erroneously classified to one of the known clusters, typically with lower probabilities, or to the outlier component (Crook et al., 2018). Secondly, for very low-abundant proteins the measured abundance distribution profiles may suffer distortion by the noise of low signals. We note, however, that many proteins even below detectable levels by Western blot or IFA were still reproducibly quantified and assigned by hyperLOPIT. Perhaps most importantly, the hyperLOPIT method reports protein steady-state locations, however there are proteins that distribute between more than one organelle and, therefore, cannot be unambiguously assigned to any one class (Crook et al., 2018; Thul et al., 2017). We found that the greatest proportion of uncertain assignments were between cytosolic and nuclear protein clusters, as well as between *Golgi*, *endomembrane vesicles*, *PM – integral*, and *PM – peripheral 2* clusters. The former is probably because many proteins shuttle between the cytosol and the nucleus, as well as that the cell disruption and fractionation method may have disrupted nuclear integrity. The latter likely reflects the intrinsic heterogeneity and dynamic nature of endomembrane compartments of the secretory pathway.

This comprehensive and unbiased spatial proteome nevertheless also opens up opportunities for discovering new, previously uncharacterized subcellular niches. The t-SNE projections indicate considerable structure in regions of these maps beyond that accessible to TAGM analysis for the lack of known markers for these clusters. The unsupervised analysis of the data (HDBSCAN) supports further genuine protein associations here. This provides routes to discover previously unrecognized cellular organization such as the *ER 2* cohort of folding-related proteins that resolved as distinct from the general ER proteome.

Pursuit of subcellular proteomes in apicomplexans has often focused on identifying molecular machinery that can elucidate the functions and mechanisms of cell compartments. Indeed, this hyperLOPIT data provides innumerable opportunities now for these important pursuits. Moreover, in combination with genetic screens, made more accessible with CRISPR/Cas9 and modern DNA sequencing methods, hyperLOPIT data provides a means to interpret the outcomes of these screens in a subcellular context. However, there is now also tremendous opportunity to understand broader processes of cell function, adaptation and evolution, by using these objective comprehensive samples of the compositional organization of the cell in combination with other systems-level data. For example, gene expression analysis reveals tight expression programs for some large molecular complexes and for invasion-related structures that likely contribute to the ordered assembly of these apicomplexan apparatuses central to pathogenicity. Further, a cell-wide view of the distribution of selective pressures and their responses, manifesting as population-level skews in gene *d*_N_/*d*_S_ and SNP-frequencies across compartments, shows how non-uniformly contemporary adaptation occurs in these parasites. Proteins changing most rapidly are enriched in rhoptries, dense granules and at the parasite surface and these likely identify molecular processes most relevant to the host-parasite arms race of attack and defense. But equally importantly, these data reveal proteins within such compartments that do not change, and that might present stable targets for therapeutic strategies against these pathogens’ core processes. Finally, deeper evolutionary questions can be asked about the relative chronology and tempo of innovation in the different cell compartments over evolutionary time. These reveal a sequence of innovation, from early to late, in functional development for: intracellular signaling cascades, metabolism, the extracellular interface, genetic networks, motility, invasion and, finally, host remodeling. Such deep insight into the stepwise chronology of the functional and proteomic innovation is without parallel in any eukaryotic system, and our analysis identifies the proteins responsible for these key events (**Table S9f**). Combating apicomplexans as pathogens requires understanding the fundamentals of their evolution as parasites, as well as the nuances of modern adaptation and molecular function. These high-resolution spatial proteomic data for apicomplexans offers a new era of discovery and advancement of our understanding and approaches to tackling these critical human pathogens.

## Supporting information

Sequences of plasmids

## ACKNOWLEDGEMENTS

We thank John Boothroyd, Peter Bradley, Mark Carrington, Vern Carruthers, Maryse Lebrun, Corinne Mercier, David Sibley, Dominque Soldati-Favre, Boris Striepen, Giel van Dooren and Gary Ward for generous gifts of antibodies used in this study. Mass spectrometry data were acquired by Mike Deery at the Cambridge Centre of Proteomics, and we thank Laurent Gatto for useful discussions. This work was supported by the Medical Research Council MR/M011690/1 to R.F.W., King Abdullah University of Science and Technology (KAUST) OSR-2015-CRG4-2610 to A.P., R.F.W. and K.S.L., Wellcome Trust Investigator Award 214298/Z/18/Z to R.F.W, a Isaac Newton Trust - Leverhulme Early Career Fellowship ECF-2015-562 to K.B, and KAUST faculty baseline funding (BAS/1/1020-01-01) to A.P.

## AUTHOR CONTRIBUTIONS

K.B., K.S.L. and R.F.W conceived this study. K.B. performed and analysed the hyperLOPIT experiments with O.M.C. and L.M.B contributing to the design and implementation of the analyses; L.K., H.K., S.B. and I.L. performed gene-tagging location validations; V.G. and E.T. the orthology analysis; T.M. and A.P. the gene expression analysis; and T.J.S. the signal peptide and transmembrane domain analysis. K.B. and R.F.W. wrote the manuscript, which was read and approved by all authors.

## METHODS

### Contact for Reagent and Resource Sharing

Further information and requests for reagents may be directed to and will be fulfilled by the corresponding author Ross F. Waller.

### Experimental Model and Subject Details

*T. gondii* tachyzoites from the strain RH and derived strains, including RH Δku80/TATi (Sheiner et al., 2011) (a kind gift from Lilach Sheiner and Boris Striepen, The University of Georgia), were maintained at 37°C with 10% CO_2_ growing in human foreskin fibroblasts (HFFs) cultured in Dulbecco’s Modified Eagle Medium supplemented with 1% heat-inactivated fetal bovine serum, 10 unit ml^−1^ penicillin and 10 μg ml^−1^ streptomycin, as described elsewhere (Roos et al., 1994). When appropriate for selection, chloramphenicol was used at 20 μM and pyrimethamine at 1 μM. Scaling up of the parasite culture for hyperLOPIT experiments was done according to the method described by Roos et al. (Roos et al., 1994).

### Method details

#### Generation of transgenic *T. gondii*

We developed a CRISPR/Cas9-assisted and PCR-mediated genomic tagging strategy to perform endogenous gene tagging with epitope tags for protein localization in *T. gondii*. It involves: 1) two-step cloning of the Cas9/sgRNA construct P5 (see **Supplementary File 3** and **Table S10a** in **Supplemental Information** for sequences and annotation of the vectors used in this study) for directing a locus-specific DNA break to facilitate homologous recombination-driven insertion of a donor DNA; 2) PCR amplification of the donor DNA fragment generating an in-frame insertion of an epitope tag as well as a drug resistance cassette amplified from one of the template vectors (P6-P10, or pPR2-HA3 (Katris et al., 2014)) using primers that include specific homology arms directing integration of this construct into the target genetic locus (**Table S10b** in **Supplemental Information**); 3) co-transfection of the plasmid generated in step 1 and the PCR product obtained in step 2 into the parasite cells.

For tagging each gene, plasmid P5 was assembled using the Golden Gate assembly method (Engler and Marillonnet, 2014). Briefly, the sgRNA was generated by PCR amplification using a gene-specific forward primer and a general reverse primer (‘Universal_sgRNA_Rv’ in **Table S10b**) from a template Golden Gate Level M plasmid P5. The resulting sgRNA containing specific protospacer sequence (PS-sgRNA) was inserted into Golden Gate Level 1 Position 2 acceptor plasmid P4 downstream of *T. gondii* U6 promoter (obtained from P1) using BsaI sites. TgU6-PS-sgRNA cassette was then combined with the ‘TgSag1 promoter – Cas9-HA-GFP – TgSag1 terminator’ cassette from P2 using BpiI to create the final plasmid P5.

For C-terminal genomic tagging, we created template plasmids containing 6xHA (P6 and P9), 3xHA (P10), or 3xV5 (P7 and P8) epitope reporters and both the dihydrofolate reductase (DHFR) and the chloramphenicol acetyltransferase (CAT) resistance cassettes (P8-P10 and P6, P7, respectively) using the Golden Gate assembly method (**Table S10a, Supplementary File 3**). For N-terminal genomic tagging with the 3xHA epitope tag, the pPR2-3HA plasmid (Katris et al., 2014) was used as a template. The resistance cassette and the reporter tag were amplified from the template plasmid using gene-specific primers that contained 3’-or 5’-end homology regions to facilitate the genomic integration by homologous recombination (**Table S10b**). Approximately 50 μg of plasmid P5 and 200 μl of the PCR reaction product containing the epitope tag, resistance gene, and the homology sequences were combined, ethanol-purified, and co-transfected into *T. gondii* RH Δku80/TATi as previously described (Heaslip et al., 2011). Parasites were selected with 1 μM pyrimethamine or 20 μM chloramphenicol. Individual clones were obtained by limiting dilution (Katris et al., 2014).

#### Immunofluorescence microscopy and immunoblotting

*T. gondii*-infected HFF monolayers grown on glass coverslips were fixed with 2% formaldehyde at room temperature for 15 min, permeabilized with 0.1% TritonX-100 for 10 min and blocked with 2% BSA for 1 h. The coverslips were then incubated with a primary antibody (see **Table S11** in **Supplemental Information** for the list of antibodies and dilutions used) for 1 h, followed by 1 h incubation with a secondary antibody (**Table S11**). Coverslips were mounted using ProLong® Diamond Antifade Mountant with DAPI (Invitrogen). Images were acquired using a Nikon Eclipse Ti widefield microscope with a Nikon objective lens (Plan APO, 100x/1.45 oil), and a Hamamatsu C11440, ORCA Flash 4.0 camera.

3D-Structured Illumination Microscopy (3D-SIM) was implemented on a DeltaVision OMX V4 Blaze (Applied Precision) with samples prepared as for widefield immunofluorescence assay (IFA) microscopy expect High Precision coverslips (Marienfeld Superior, No1.5H with a thickness of 170 μm ± 5 μm) were used in cell culture and Vectashield (Vector Laboratories) was used as mounting reagent. Samples were excited using 405, 488 and 594 nm lasers and imaged with a 60x oil immersion lens (1.42 NA). The structured illumination images were reconstructed in softWoRx software version 6.1.3 (Applied Precision). All fluorescence images were processed using Image J software (http://rsbweb.nih.gov./ij/).

For immunoblotting, performed during optimization of cell disruption and density gradient fractionation, approximately 5-10 × 10^7^ gene-tagged parasites were purified from the host cell debris by filtration through 3-µm-pore-size polycarbonate film membrane filters (Nuclepore Track-Etch Membrane, Whatman) and collected and washed in PBS by centrifugation at 1,700 × *g*_max_ for 10 min at room temperature. The cell pellets were directly resuspended to an equivalent number density of approximately 5 × 108 ml^−1^ in NuPage LDS Sample Buffer (Thermo Fisher Scientific) supplemented with dithiothreitol (DTT) to a final concentration of 50 mM and incubated at 70°C for 10 min to extract, reduce, and denature proteins. For the hyperLOPIT density gradient assessment, aliquots of the gradient fractions containing 0.5 µg total protein were prepared in NuPAGE LDS Sample Buffer as described above. Proteins were resolved by SDS-PAGE using NuPAGE 4-12% Bis-Tris Protein Gels (Thermo Fisher Scientific) and electrotransferred onto 0.2-µm-pore-size nitrocellulose membranes (Amersham Protran Supported, GE Healthcare) using either XCell SureLock Mini-Cell with XCell II Blot Module or Mini Gel Tank with Blot Module (Thermo Fisher Scientific) according to the manufacturer’s instructions. The membranes were blocked in 5% (w/v) non-fat dry milk in tris-buffered saline solution containing 0.05% (w/v) of Tween 20 (TBST) and probed with primary and secondary antibodies (**Table S11**). Protein bands were visualized via chemiluminescence detection using SuperSignal West Pico Chemiluminescent Substrate (Thermo Scientific).

#### Sample preparation for hyperLOPIT

Approximately 10^10^ (**Table S1** in **Supplemental Information**) freshly egressed extracellular tachyzoites were purified from the host cell debris by filtration through 3-µm-pore-size polycarbonate film membrane filters (Nuclepore Track-Etch Membrane, Whatman). The cells were washed with chilled PBS (pH 7.4) three times by centrifugation at 3000 × *g*_max_, 4°C and resuspended to a final cell density of 5 × 10^8^ ml^−1^ in a chilled homogenization medium (HB: 0.25 M sucrose, 10 mM HEPES•KOH pH 7.4, 1 mM EDTA) supplemented with proteinase inhibitors (cOmplete™ EDTA-free Proteinase inhibitor cocktail, Roche).

The cells were mechanically lysed by nitrogen cavitation (Hunter and Commerford, 1961; Simpson, 2010) using a Parr Instruments cell disruption vessel model 4639 (45 ml volume) at 2,000 PSI (approximately 138 bar). The system with cell suspension was allowed to equilibrate on ice for 15 min with occasional gentle agitation. The content was discharged from the vessel through the release valve at a flow rate of approximately two droplets per second. Differential centrifugation was used to return intact and poorly dispersed cell material to a subsequent cavitation cycle. The unlysed material was removed by centrifugation as described in **Table S1**. The resulting supernatant was considered the cell homogenate. In some cases (**Table S1**), the homogenate was treated with 500 U of the nuclease Benzonase (Sigma-Aldrich) for 20 min at room temperature and for a further 10 min at 4°C in order for its viscosity to be reduced.

#### HyperLOPIT subcellular fractionation

The suspension of membrane vesicles and subcellular particles was resolved on an iodixanol density gradient as described in (Christoforou et al., 2016; Mulvey et al., 2017). Briefly, crude subcellular particles were enriched by ultracentrifugation of the homogenate underlaid with 6 and 25% (w/v) iodixanol solutions in HB for 1.5 h at 100,000 × *g*_max_, 4°C (SW32Ti rotor, Optima L-80XP ultracentrifuge, Beckman) with the maximum acceleration and minimum deceleration. An aliquot of the supernatant enriched with cytosolic and soluble proteins was taken and mixed with six volumes of acetone chilled to −20°C and removed to −20°C to precipitate proteins from the solution. Opaque bands at the interfaces of the iodixanol layers containing enriched subcellular membranes and particles were collected, diluted with HB to bring the iodixanol concentration below 6% (w/v), and pelleted from residual soluble proteins by ultracentrifugation for 1 h at 200,000 × *g*_max_, 4°C (SW55Ti rotor, Beckman). The pellets were resuspended in 25% (w/v) iodixanol in HB using a Dounce tissue grinder (max. volume 2 ml, Kimble, pestle A clearance 0.0030-0.0050 in., pestle B clearance 0.0005-0.0025 in.) and underlaid beneath a linear pre-formed density gradient (equal volumes of 8, 12, 16, and 18% (w/v) iodixanol solutions in HB allowed to diffuse at 4°C overnight). The sample was centrifuged for 8 h at 100,000 × g_max_, 4°C (VTi65.1 rotor, Beckman) with the maximum acceleration and minimum deceleration allowing for isopycnic separation of subcellular particles and membranes. The resolving gradient was harvested into 23 approximately equal-volume fractions by piercing the ultracentrifugation tube bottom and allowing the liquid to dispense dropwise under gravity flow. Aliquots were taken from each fraction to determine the average density through measuring the refractive index (Eclipse Handheld Refractometer 45-02, sugar 0-32%, Billingham and Stanley), and for protein concentration assessment by the BCA protein assay (Thermo Fischer Scientific) according to the manufacturer’s instructions. The distribution of several known organelle marker proteins in the gradient fractions was assessed by Western blotting using aliquots containing 0.5 µg total protein.

#### Protein extraction and proteomic sample generation

In experiments Toxoplasma LOPIT 1 (TL1) and TL3, the harvested fractions of the density gradient were stored at −80°C until used; proteins were extracted from the gradient fractions by precipitation with 10% (w/v) trichloroacetic acid (TCA) as described elsewhere (Link and Labaer, 2011). In experiment TL2, each fraction of the gradient was diluted with 0.8 ml HB and centrifuged for 1 h at 100,000 × *g*_max_, 4°C (TLA-55 rotor, Optima MAX-XP benchtop ultracentrifuge, Beckman). The supernatant was carefully aspirated and discarded, membrane pellets were resuspended in 0.8 ml HB by repeated tube inversion and pelleted again by ultracentrifugation. The supernatant was discarded, and the resulting membrane-enriched pellets were stored at −80°C until used. Protein (TL1 and TL3) or membrane (TL2) pellets, including the acetone-precipitated proteins from the cytosol-enriched fraction, were resolubilized in triethylammonium bicarbonate (TEAB) buffered solution (pH 8.3) containing either 0.1% SDS (TL1) or 8 M urea, 0.2% SDS (TL2 and 3) assisted by sonication (5 cycles of 30 s ON, 30 s OFF at high power, Bioruptor Plus ultrasonic disintegrator, Diagenode). Protein concentration was measured by the BCA assay.

Sequential gradient fractions were aggregated to nine pools containing 60 to 100 µg protein and maximizing distinct subcellular marker protein distributions based on Western blots analysis (see pooling strategies in **Table S1**). A tenth fraction was derived from the soluble protein-containing fraction. Proteins were reduced with 10 mM Tris(2-carboxyethyl)phosphine (TCEP; Sigma-Aldrich) for 1 h at room temperature followed by alkylation of cysteine residue side chain thiol groups with iodoacetamide (Sigma-Aldrich) at approximately 17 mM final concentration for 30 min at room temperature in the dark. Six volumes of pre-chilled (−20°C) acetone were added to the reaction mixtures and proteins were allowed to precipitate overnight at −20°C. The samples were centrifuged at 16,000 × *g*_max_ for 10 min at 4°C, the supernatant was carefully aspirated and discarded, and the protein pellets were air-dried at room temperature for 5 min.

Acetone-precipitated protein pellets were resuspended in 100 mM TEAB-buffered solution (pH 8.3) with the assistance of sonication (Bioruptor Plus, Diagenode, 5 cycles of 30 s ON, 30 s OFF, high power) and digested with 1 µg of sequencing-grade trypsin (Promega) for 2 h at 37°C followed by the addition of another 1 µg aliquot of the enzyme and incubation at 37°C overnight. The digests were centrifuged for 10 min at 16,000 × *g*_max_ at 4°C to remove any insoluble material, and the supernatants were transferred to new 1.5 ml Protein LoBind microcentrifuge tube (Eppendorf) and labelled with TMT10plex isobaric tagging reagents (Thermo Fisher Scientific) according to the manufacturer’s instructions. Briefly, 0.8 mg of TMT10plex reagents were brought to room temperature and dissolved in 41 µl of LCMS-grade acetonitrile immediately before use. The peptide digest samples (approximately 100 µl) were transferred to the TMT10plex reagent vials and the reaction mixtures were incubated at room temperature for 1-2 h with constant agitation (800 RPM, PHMT thermomixer, Grant Bio Instruments). The reaction was stopped by adding 8 µl of 5% (v/v) hydroxylamine solution and incubation for 15 min at room temperature with agitation. The TMT-labelled fractions were combined and reduced to dryness in a refrigerated (4°C) vacuum centrifuge (Labconco).

The combined TMT-labelled peptide samples were desalted using C18 solid-phase extraction (SPE) cartridges (SepPak C18, 100 mg sorbent, Waters). The dry samples were resuspended in 0.8 ml of 0.5% (v/v) trifluoroacetic acid (TFA) solution in HPLC-grade water with the assistance of sonication (Bioruptor Plus, Diagenode, 5 cycles of 30 s ON, 30 s OFF, high power). The SPE resin was conditioned with 1.6 ml of LCMS-grade acetonitrile and equilibrated in 0.1% (v/v) aqueous TFA solution (a total volume of 1.6 ml). The peptide samples were loaded onto the cartridges under the gravity-flow. The cartridges were washed with 1.6 ml of 0.1% (v/v) aqueous TFA solution to remove salts and other polar low-molecular-weight contaminants and equilibrated in 0.5% (v/v) aqueous solution of acetic acid (a total volume of 1.6 ml). The peptides were eluted from the resin using 1.6 ml of 70% (v/v) LCMS-grade acetonitrile, 0.5% (v/v) acetic acid solution in HPLC-grade water and reduced to dryness in a refrigerated (4°C) vacuum centrifuge (Labconco).

#### High-pH reversed-phase fractionation of peptides

The TMT10plex-labelled desalted peptide samples were fractionated by high-pH reverse-phase chromatography on an Acquity UPLC BEH C18 column (2.1-mm i.d. × 150-mm; 1.7-μm particle size) with a VanGuard pre-column (2.1 × 5 mm) packed with the same resin (both from Waters) using an Acquity UPLC system equipped with an autosampler, a binary solvent manager, and a diode array detector (Waters). The following solutions for gradient elution were used: 20 mM ammonium formate in HPLC-grade water, pH 10 (Eluent A); 20 mM ammonium formate in LCMS-grade acetonitrile: HPLC-grade water 80:20 (v/v), pH 10 (Eluent B).

The dried peptide samples were resuspended in 100 µL of 5% (v/v) Eluent B in Eluent A, sonicated (5 cycles of 30 s ON, 30 s OFF, high power, Bioruptor Plus, Diagenode), spun for 10 min at 16,000 × *g*_max_ to remove any insoluble material, and the supernatants were injected onto the column equilibrated with at least 20 column volumes of 95% Eluent A: 5% Eluent B. A flow rate of 0.244 ml min^−1^ was maintained. The percentage of Eluent B was varied according to the following program: 5% for 10 min, 5 to 75% over 50 min, a ramp to 100% over 2 min followed by 5.5 min at 100%, switching to 5% and equilibration for 10 min. Fifty 1-min fractions were collected along the elution profile of the peptides (approximately from minute 10 to 60 of the program) and reduced to dryness. For the downstream LC-MS analysis, the fractions corresponding to each TMT10plex set were concatenated into 15-18 samples by combining pairs of fractions which eluted at different time points during the gradient, e.g., fraction 1, 16, and 31, fraction 2, 17, and 32, etc.

#### LC-MS analysis of peptides

All mass spectrometry analyses were performed on an Orbitrap Fusion™ Lumos™ Tribrid™ instrument coupled to a Dionex Ultimate™ 3000 RSLCnano system (Thermo Fisher Scientific) as described in (Geladaki et al., 2019).

Briefly, each of the fractionated samples was resuspended in 30 μL of 0.1% (v/v) aqueous solution of formic acid. Approximately 1 μg of peptides was loaded per injection for LC-MS/MS analysis.

The nano-flow liquid chromatography method for LC-MS/MS was set as follows. Eluent A was 0.1% (v/v) formic acid solution in water. Eluent B was 80% (v/v) aqueous acetonitrile supplemented with formic acid to a final concentration of 0.1% (v/v). The sample loading solvent was 0.1% (v/v) formic acid in water. All solvents and reagents were of HPLC gradient grade or better. Peptides were loaded onto a micro precolumn (300 μm i.d. × 5 mm, particles were C18 PepMap 100, 5 μm particle size, 100 Å pore size, Thermo Fisher Scientific) using the loading pump for 3 min. After this, the valve was switched from load to inject. Peptides were separated on a Proxeon EASY-Spray column (PepMap RSLC C18, 50 cm × 75 μm i.d., 2 μm particle size, 100 Å pore size, Thermo Fisher Scientific) using a 2-40% (v/v) gradient of acetonitrile supplemented with 0.1% (v/v) formic acid at 300 nL min^−1^ over 93 min. A wash step (90% Eluent B for 5 min) was included, followed by re-equilibration into Eluent A. The total run time was 120 min.

The MS workflow parameters were set as follows using the Method Editor in XCalibur v3.0.63 (Thermo Fisher Scientific) for the SPS-MS3 acquisition method. Detector type: Orbitrap; Resolution: 120,000; Mass range: Normal; Use quadrupole isolation: Yes; Scan range: 380-1,500; RF lens: 30%; AGC target: 4e5; Max inject time: 50 ms; Microscans: 1; Data type: Profile; Polarity: Positive; Monoisotopic peak determination: Peptide; Relax restrictions when too few precursors are found: Yes; Include charge state(s): 2-7; Exclude after n times: 1; Exclusion duration (s): 70; Mass tolerance (p.p.m.): Low: 10; high: 10; Exclude isotopes: Yes; Perform dependent scan on single charge state per precursor only: Yes; Intensity threshold: 5.0e3; Data-dependent mode: Top speed; Number of scan event types: 1; Scan event type 1: No condition; MSn level: 2; Isolation mode: Quadrupole; Isolation window (*m/z*): 0.7; Activation type: CID; CID collision energy (%): 35; Activation Q: 0.25; Detector type: Ion trap; Scan range mode: Auto; *m/z*: Normal; Ion trap scan rate: Turbo; AGC target: 1.0e4; Max inject time (ms): 50; Microscans: 1; Data type: Centroid; Mass range: 400-1200; Exclusion mass width: *m/z*: Low: 18; high: 5; Reagent: TMT; Precursor priority: Most intense; Scan event type 1: No condition; Synchronous precursor selection: Yes; Number of precursors: 10; MS isolation window: 0.7; Activation type: HCD; HCD collision energy (%): 65; Detector type: Orbitrap; Scan range mode: Define *m/z* range; Orbitrap resolution: 60,000; Scan range (*m/z*): 100-500; AGC target: 1.0e5; Max inject time (ms): 120; Microscans: 1; Data type: Profile; AGC, automatic gain control; HCD, higher-energy collisional dissociation; CID, collision-induced dissociation.

An electrospray voltage of 2.1 kV was applied to the eluent via the electrode of the EASY-Spray column. The mass spectrometer was operated in positive ion data-dependent mode for SPS-MS3. The total run time was 120 min.

### Quantification and statistical analysis

#### Raw LC-MS data processing and quantification

Raw LC-MS data files were processed with Proteome Discoverer v2.1 (Thermo Fisher Scientific) using the Mascot server v2.6.0 (Matrix Science). The annotated protein sequences for *T. gondii* strain ME49 retrieved from the ToxoDB.org database (release 29, downloaded on 12.10.2016) was used along with common contaminants from the common Repository of Adventitious Proteins (cRAP) v1.0 (112 sequences, adapted from the Global Proteome Machine repository, https://www.thegpm.org/crap/). Precursor and fragment mass tolerances were set to 10 ppm and 0.8 Da, respectively. Trypsin was set as the enzyme of choice and a maximum of 2 missed cleavages were allowed. Static modifications were carbamidomethyl (C), TMT6plex (N-term), and TMT6plex (K). Dynamic modifications were oxidation (M), deamidated (NQ), TMT6plex (S/T). Percolator version 2.05 (Kall et al., 2008; Käll et al., 2007, 2008) was used to assess the false discovery rate (FDR) and only high-confidence peptides were retained.

Quantification at the MS3 level was performed within the Proteome Discoverer workflow using the Most Confident Centroid method for peak integration and integration tolerance of 20 p.p.m. An isolation interference threshold was set to 50%. Reporter ion intensities were adjusted to correct for the isotopic impurities of the different TMT reagents (manufacturer’s specifications).

Protein grouping was carried out according to the strict parsimony principle. Only proteins with a full reporter ion series and medium (q ≤ 0.05) or high (q ≤ 0.01) FDR confidence level were retained. Non-*Toxoplasma* proteins were removed for downstream analysis.

#### Protein subcellular localization prediction by machine learning methods

Data analysis was performed using the R (R Core Team, 2018) Bioconductor (Gentleman et al., 2004) packages MSnbase v2.8.3 (Gatto and Lilley, 2012) and pRoloc v1.22.1 (Gatto et al., 2014) as described in (Breckels et al., 2016; Crook et al., 2019). Briefly, the quantitative proteomics datasets obtained in three independent hyperLOPIT experiments were subset for shared proteins with full TMT10plex quantitation data series, thereby yielding a concatenated dataset of 3,832 features (proteins) quantified across 30 samples (TMT10plex intensity values). The raw quantitation values output from Proteome Discoverer were normalized feature-wise to the sums of intensities across samples followed by variance-stabilizing normalization (Huber et al., 2002). Principle component analysis (PCA) and t-distributed Stochastic Neighbour Embedding (t-SNE) (van der Maaten et al., 2008) were used for dimensionality reduction and data visualization.

For t-SNE, the data were preprocessed as follows. First, the normalized data were centered, scaled, and PCA-transformed. The top principle components accounting for a cumulative variance of approximately 99% were filtered and subjected to a statistical whitening transformation such that each principle component had variance 1. An embedding of the preprocessed data into two dimensions was produced using a perplexity of 50 and exact gradient calculation for a maximum of 10,000 iterations. The computed coordinates were recorded and used to obtain the two-dimensional data projection shown throughout the text.

For the Bayesian machine-learning classification of protein locations, a set of 718 manually curated marker proteins defining 26 subcellular classes was compiled using previously published data (656 proteins) and in-house protein localization by epitope-tagging and immunofluorescence microscopy (62 proteins). A Bayesian generative classifier based on t-augmented Gaussian mixture models (TAGM) was used to probabilistically attribute proteins marked as ‘*unknown*’ to the 26 classes defined by the marker set using maximum *a posteriori* prediction (TAGM-MAP) or Markov-chain Monte-Carlo (TAGM-MCMC) methods as described in (Crook et al., 2019). TAGM-MAP was used to obtain maximum *a posteriori* probability of each protein to belong to one of the 26 classes or to an outlier component. We determined the model parameters by performing 100 iterations of the expectation-maximization (EM) algorithm using the default priors and confirmed convergence by assessing the log-posterior plot. The class with the highest TAGM-MAP allocation probability was defined as the most likely protein subcellular location. To retain only high-confidence assignments, a threshold of 99% was set on the posterior localization probability, which was defined as a product of the allocation probability and the complement of the outlier probability: *p*_*localisation*_ = *p*_*allocation*_ ⋅ (1 − *p*_*outlier*_). To quantify uncertainty in the allocation of proteins to organelles, we applied TAGM-MCMC. To obtain samples from the posterior localization distributions, we performed inference in this model using Markov-chain Monte-Carlo (Gilks et al., 1995). The collapsed Gibbs sampler was run in parallel for 9 chains, with each chain run for 25,000 iterations. We discarded 10,000 iterations for burn-in and thinned the chain by retaining every 20th sample. For the combined 3 replicate experiment, we discarded 5 chains because they were deemed not to have converged from visual inspection. We used the Gelman-Rubin’s diagnostic (Gelman and Rubin, 1992) to further asses convergence of the remaining Markov chains. We computed a potential scale reduction factor 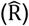) of 1.02, which is less than 1.2, upon which we concluded convergence of our algorithm. The 750 samples from each of these 4 chains are then pooled together for further downstream processing.

#### Prediction of signal peptides and transmembrane domains

SignalP 5.0 (Almagro Armenteros et al., 2019) was used to predict cleaved signal peptides in *T. gondii* ME49 annotated protein sequences with organism group set to Eukarya.

Transmembrane (TM) span prediction was performed using TMHMM 2.0 (Krogh et al., 2001). It is known signal peptides can create TM false positives, so the results of TMHMM and SignalP analyses were compared and whenever a protein was predicted to have an N-terminal TM span and simultaneously a signal peptide the first TM span predicted by TMHMM was removed (**Table S3**).

#### Subcellular distribution of quantitative gene and protein characteristics

The TAGM-MAP-predicted subcellular proteomes (localization probability > 99%) were used to quantitatively assess the distributions of the calculated protein pI (**Table S7a**), the protein-averaged ratio of non-synonymous to synonymous mutation rates (*d*_N_/*d*_S_) (Lorenzi et al., 2016) (**Table S7b**), the density of single nucleotide polymorphism (SNP density; data retrieved from the ToxoDB.org; **Table S7c**), and the average phenotype score determined by Sidik et al. in the whole-genome CRISPR/Cas9-mediated loss-of-fitness screen in *T. gondii* (Sidik et al., 2016) (**Table S7d**). These analyses sought to identify subcellular proteomes whose distributions of these quantities are unusual given the spatial proteome map described here. No prior knowledge of the true distributions of these quantities was available, hence, a non-parametric significance test based on an exact inference of null distributions of class-specific means through random permutation of class labels was performed.

In brief, for each of the analyzed quantities, the vector of organelle class labels was randomly permuted with replacement, and the mean value was computed for each subcellular class. The procedure was repeated *m* = 106 times to infer the null distributions of class means. To test the hypothesis that the observed distribution of the quantity could emerge by chance, the number *b* of instances from the null distribution at least as extreme as the observed mean was counted. The approximate *p*-value was then calculated according to the following formula: *p* = (*b* + 1)/(*m* + 1) (Phipson and Smyth, 2010; Young et al., 2003). The resulting *p*-values were adjusted for multiple comparison using the Benjamini-Hochberg method (Benjamini and Hochberg, 1995).

#### Analysis of gene co-expression

RNA-Seq gene expression data sets were collected from ToxoDB (**Table S6**). All data was downloaded as FPKM values. In parallel, the combined data sets were treated in three different ways: i) non-normalized, ii) z-transformed, and iii) quantile normalized. Quantile normalization was done using the ‘normalize.quantiles’ function in the R package preprocessCore (https://github.com/bmbolstad/preprocessCore). Co-expression levels of genes across the data sets were calculated as both Pearson and Spearman rank correlations.

To test if genes within a given cluster showed signs of co-expression, all pair-wise co-expression values between members of the cluster were compared to the co-expression between members of the cluster and all genes outside the cluster. The distributions of co-expression levels were compared using the Mann-Whitney *U* test of medians, and Cohen’s *d* test of effect size. All tests were carried out in R (R Core Team, 2018). All normalization procedures and correlation measures yielded highly similar results (**Table S6**).

#### Gene orthology analysis

Eukaryote-wide protein orthogroups were defined using OrthoFinder (Emms and Kelly, 2015) between 79 proteomes spanning 12 levels of evolutionary divergence (**Table S9a**) using default OrthoFinder parameters. Either predicted or reviewed proteomes were downloaded from protein databases UniProt (The UniProt Consortium, 2019), ToxoDB (Gajria et al., 2007), PlasmoDB (Aurrecoechea et al., 2009), PiroplasmaDB (Aurrecoechea et al., 2010), and CryptoDB (Heiges, 2006). Proteomes were compared in all vs. all searches using Diamond (Buchfink et al., 2014), and *T. gondii* protein orthologues were inferred under stringent criteria (reciprocal best hits, RBHs) in all 78 species (Emms and Kelly, 2015). A binary bit string representing the absence/presence profile of each *T. gondii* protein orthologue at all evolutionary levels was computed from the OrthoFinder output (**Table S9c**). A conservation score for each *T. gondii* protein against a predefined set binary conservation profiles (**Table S9d**) was computed using pairwise Jaccard index. Based on the highest conservation score, each *T. gondii* protein was assigned a conservation profile (**Table S9e**).

To test for evidence of cell compartments enriched in new orthogroups at a given phylogenetic position, protein sets for each conservation profile were then tested for enrichment across all of the hyperLOPIT-derived annotation classes and a *p*-value for the likelihood of a given enrichment to have occurred by chance was obtained using a hypergeometric test (**Table S9f**).

#### Signal peptide and membrane-spanning sequence properties

Analyses of signal peptide (SP) and transmembrane span sequence properties were performed as previously described (Parsons et al., 2019) with only minor modifications. SP-containing proteins and monotopic integral membrane proteins were identified using SignalP 4.1 (Petersen et al., 2011) and TMHMM 2.0 as described above. Phobius (Käll et al., 2004) was used to estimate the initial TM span edge positions and the cytoplasm-exoplasm transmembrane topology.

*T. gondii* protein sequences were augmented with sequence information from close apicomplexan homologues identified in our OrthoFinder-based search for gene homologues across 79 eukaryotic taxa (see above). Resulting family groups all had a single, consistent organelle or subcompartment annotation that was derived from the *T. gondii* query protein.

Families of sequences were multiply aligned using Clustal Omega (Sievers and Higgins, 2014) with default parameters. TM span edge positions were refined using the multiple alignment of each homologue family. First, the edges of the TM span (initially predicted by Phobius) were adjusted within a region of five residues either side by selecting the point in the alignment with the maximum difference in GES-scale (Engelman et al., 1986) hydrophobicity (summed over all proteins in the alignment) between the adjacent five residues on the side of the TM span and the adjacent five residues on the opposite side. Next, the edge positions were trimmed or extended according to the average hydrophobicity over the whole alignment. If the mean hydrophobicity of the next residue exceeded 1.0 kcal mol^−1^ (Gly or more hydrophobic), the edge was extended. Similarly, if the mean hydrophobicity of an edge residue was below 1.0 kcal mol^−1^, the edge was trimmed. Finally, individual protein adjustments were made, extending or trimming positions for each span sequence. Accordingly, individual TM span edges were trimmed if they ended in a gap or a hydrophilic residue (defined here as Arg, Lys, Asp, Glu, Gln, Asn, His, or Ser) or extended if the next residue was suitably hydrophobic (Phe, Met, Ile, Leu, Val, Cys, Trp, Ala, Thr, or Gly).

Next, families of proteins were multiply aligned again using Clustal Omega, and the following additional checks were made for a comparable TM span, comparing each OrthoFinder hit with the query: (1) the length of the protein must not differ by more than 200 residues; (2) there must not be more than four gap insertions in the TM span region; (3) the separation from the TM span to the N-terminus must not differ by more than 75 residues; and (4) there must be a cursory similarity between span sequences (mean, aligned regional BLOSUM62 score > 0.8).

SP cleavage site positions were taken from the SignalP predictions.

The resulting sequence counts across compartment-specific protein families are given in **Table S8a** and **Table S8c**.

Given that families contain different numbers of protein sequences with different degrees of similarity, each protein was weighted according to its dissimilarity to all other sequences in the whole data set. Dissimilarity weights for each protein (*w*_*p*_) were obtained using a BLAST+ search of each sequence (maximum e-value 10^−20^) against a database of all the protein sequences and were calculated as:

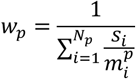

Here, *s*_*i*_ is the BLAST+ bit score of the aligned high-scoring database hit *i* (from a total of *N*_*p*_ hits) and 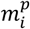 is the maximum possible bit score value, the bit score if the query were compared with itself over the same alignment region. Accordingly, the dissimilarity weight is 1.0 if the search only finds itself and approximately 1/*N* if it finds *N* very similar sequences. This protects against large and/or well-conserved protein families having an undue influence on the measurement of general TM span or SP properties.

The distributions of TM span length of compartment-specific protein families were compared pairwise using the Mann-Whitney *U* test. The resulting *p*-values (smaller means less similarity between the distributions) were used to cluster the distributions by Ward’s method (Figure 5*B*).

The positional abundance of amino acid residue types in SP sequences (**Table S8b**) was analyzed for the region of 15 residues [-10, +5] anchored on the signal cleavage site using Fisher’s exact test on the counts of each residue type (vs. other types) in a query compartment-specific protein set compared to all other protein sets. All counts have been corrected for familial similarity using the measure of uniqueness described above and re-scaling for the original protein count.

#### Protein Sequence Logo Plots

The frequency of residue occurrence in SPs and flanking regions of *T. gondii* and their close homologues was visualized using logo plots. Logo plots were generated by specially written Python scripts (available at github.com/tjs23/logo_plot), after randomly sampling 1000 sequences for each data set, from position-specific residue abundance probabilities calculated from dissimilarity-weighted sequences. Sampling sequences using dissimilarity weights (as defined above) reduced the effect of similar, redundant sequences and allowed better comparison of groups containing differently sized homologous protein families. Different proteins within each subgroup were aligned by anchoring their sequences at the SP cleavage site, prior to the generation of logo plots (Figures 5*A* and **S6**).

#### Data and software availability

The mass-spectrometry-based proteomics data have been deposited to the ProteomeXchange Consortium (Deutsch et al., 2017) via the PRIDE (Perez-Riverol et al., 2019) partner repository with the dataset identifier PXD015269 and 10.6019/PXD015269. The data are integrated into ToxoDB (https://toxodb.org/toxo/app/record/dataset/DS_eda79f81b5) (Gajria et al., 2007). The protein-level dataset generated in this study is available in the R Bioconductor pRolocdata package (version ≥ 1.25.2). An interactive interface to the annotated spatial proteome data is available through the pRolocGUI application (version 1.18.0) or via a web-based R Shiny application at https://proteome.shinyapps.io/toxolopittzex/.

## SUPPLEMENTARY FIGURE TITLES AND LEGENDS

**Figure S1. Abundance distribution profiles of marker and unknown proteins measured across three hyperLOPIT experiments, related to Figure 1.**

***A.*** The normalized intensities are shown on the Y-axis (Abundance). Data from three 10plex hyperLOPIT experiments are concatenated to yield a single 30plex dataset. The fractions are labelled according to the TMT10plex tag used for labelling.

***B.*** A dendrogram of hierarchical clustering of marker protein abundance distribution profiles. For each subcellular class, a consensus abundance distribution profile was generated by averaging the profiles of the respective marker proteins. Hierarchical clustering was performed using Euclidean distance and unweighted pair group method with arithmetic mean (UPGMA).

**Figure S2. Validation of hyperLOPIT-predicted subcellular locations of select uncharacterized proteins by epitope tagging and immunofluorescence microscopy, continued from Figure 2.**

Proteins belonging to the *apical 1* and *2* (***A***), *rhoptries 1* and *2* (***B***), *micronemes* (***C***), *dense granules* (***D***), *Golgi* (***E***), *PM – peripheral 2* (***F***), *mitochondrion-membranes* and -*soluble* (***G***), and *apicoplast* (***H***) clusters are grouped in panels. Scale bar = 10 µm for all.

**Figure S3. The posterior localization probabilities of 3,832 *T. gondii* proteins determined by a supervised Bayesian classification method TAGM-MAP, related to Figure 3.**

Proteins are grouped by the most probable subcellular class, as per the TAGM-MAP classification result, and ranked on the x-axis by their localization probability. The marker proteins are shown in cyan, and the allocated proteins are in red. For each protein, the probability to belong to the outlier component is also shown in grey. The vertical dotted line in each panel indicates the protein localization prediction cutoff (localization probability threshold of 0.99). Only proteins with the localization probability above the threshold of 0.99, i.e. with the rank below the cutoff, retained their class label, whereas the rest of the proteins were labelled as ‘*unassigned*’.

**Figure S4. Distributions of select protein sequence features and properties in the spatial proteome of *T. gondii* extracellular tachyzoite, related to Figures 3, and 5.**

***A.*** A t-SNE projection of the 30plex hyperLOPIT data on 3,832 *T. gondii* proteins with monotopic (blue) and polytopic (red) integral membrane proteins highlighted. TMHMM 2.0 was used to predict transmembrane (TM) spans. The TM spans that overlapped with the signal peptide predicted by SignalP were removed.

***B.*** Same as in ***A*** but with the proteins predicted to have a signal peptide (SignalP 5.0) highlighted in green.

***C.*** Same as in ***A*** but showing the distribution of protein charge. Proteins are colored according to protein pI computed based on the amino acid sequence. The scale is from red for acidic proteins to blue for basic proteins with the midpoint at pI = 7.4 (colorbar in the bottom-right corner of the panel).

**Figure S5. Gene co-expression levels for 26 gene clusters corresponding to *T. gondii* extracellular tachyzoite subcellular compartments determined by hyperLOPIT, related to Figure 4.**

The co-expression levels for the genes within hyperLOPIT-defined cluster are shown as light-blue bars in histograms. The co-expression levels between the cluster members and all the genes that are not members of the cluster are shown as orange bars. The Y-axis shows the fraction of gene pairs. The X-axis shows Pearson correlation (the range is from −1 to 1) of non-normalised quantitative transcriptomics data retrieved from ToxoDB.org. Cohen’s *d* values with *effect size* descriptors are shown above each plot.

**Figure S6. Logo plots of signal peptide (SP) sequences for select cohorts of *T. gondii* proteins, related to Figure 6.**

The logo plots show positional abundances of amino acid residue types within and immediately downstream of the SP cleavage site. Proteins from hyperLOPIT-defined *T. gondii* compartment-specific sets and their close homologues from Apicomplexa were aligned anchored at the SP cleavage site (positon 0). Logo plots were generated after randomly sampling 1000 sequences for each data set from position-specific residue abundance probabilities calculated from dissimilarity-weighted sequences. The plots were generated in the same way as in Figure 5*A*.

**Figure S7. Mapping of the average CRISPR/Cas9-mediated gene knockout phenotype score (A), evolutionary selective pressure (B), and genetic polymorphism (C) on the 30plex hyperLOPIT t-SNE projection of *T. gondii* extracellular tachyzoite spatial proteome data, related to Figure 6.**

***A.*** Distribution of the average CRISPR/Cas9-mediated gene knockout phenotype score (Sidik et al., 2016). The range is from blue for essential genes to yellow for dispensable genes with the midpoint set at −2.4.

***B.*** Distribution of the protein-average ratio of non-synonymous to synonymous point mutations (*d*_N_/*d*_S_) (Lorenzi et al., 2016). The scale is clipped at the 99-% quantile of the *d*_N_/*d*_S_ range. Data points with extremely high (top-1%) *d*_N_/*d*_S_ values are colored in yellow.

***C.*** Distribution of the density of single nucleotide polymorphism (SNP) per Kb of protein-coding sequence (CDS) of genes. The SNP density data were retrieved from ToxoDB.org. As in ***A***, the scale is clipped at the 99-% quantile of the data range, with extremely high values (top-1%) shown in yellow.

## Supplemental Information

Supplementary File 1: Supplementary Figures S1-S7

Supplementary File 2: Supplementary Tables S1-S11

Supplementary File 3: sequences of plasmids in FASTA format

