## Supplementary Figures S1-S7 for "A subcellular atlas of *Toxoplasma* reveals the functional context of the proteome"

Figure S1

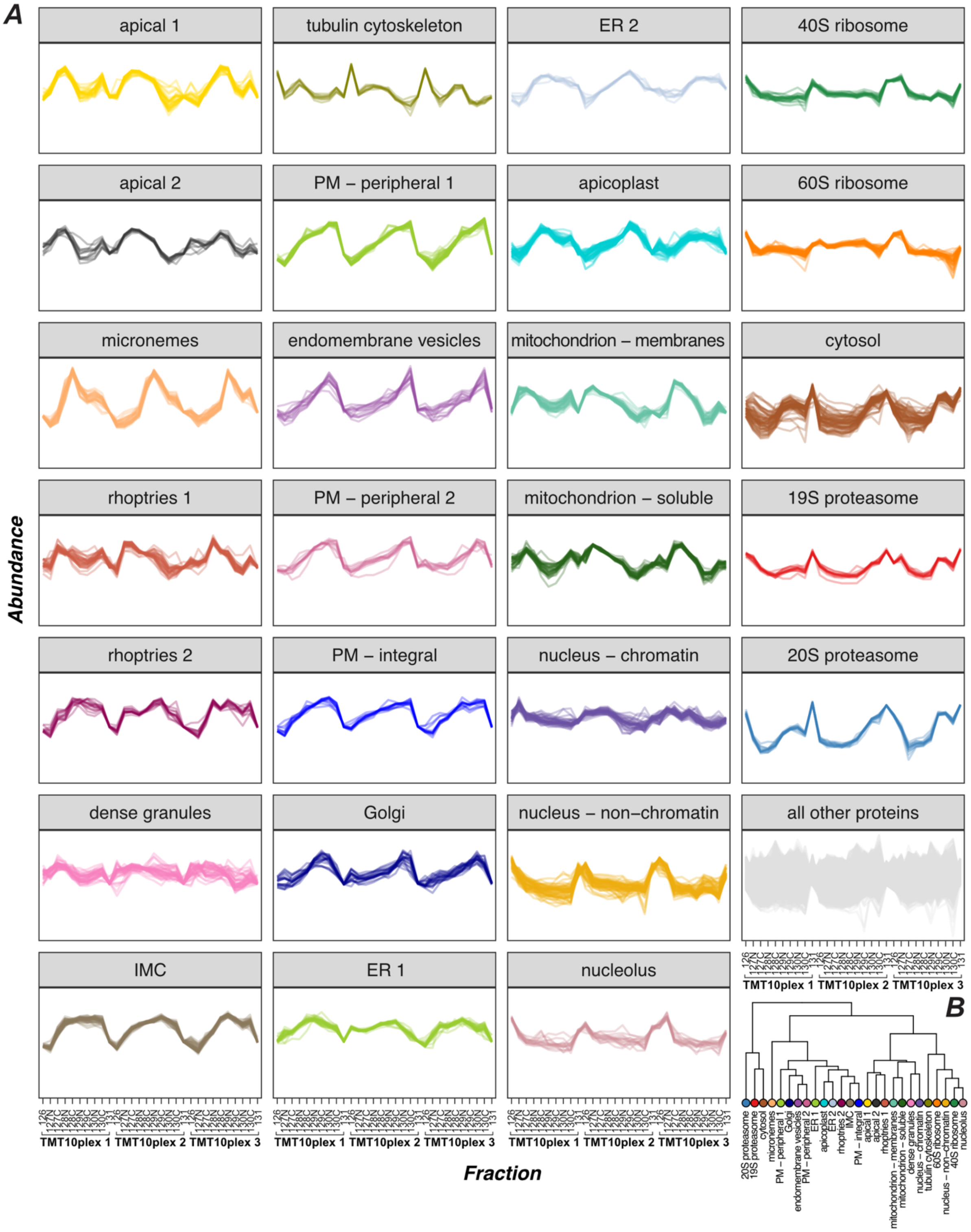


Figure S1. Abundance distribution profiles of marker and unknown proteins measured across three hyperLOPIT experiments, related to Figure 1.

***A***. The normalized intensities are shown on the Y-axis (Abundance). Data from three 10plex hyperLOPIT experiments are concatenated to yield a single 30plex dataset. The fractions are labelled according to the TMT10plex tag used for labelling.

***B.*** A dendrogram of hierarchical clustering of marker protein abundance distribution profiles. For each subcellular class, a consensus abundance distribution profile was generated by averaging the profiles of the respective marker proteins. Hierarchical clustering was performed using Euclidean distance and unweighted pair group method with arithmetic mean (UPGMA).

Figure S2

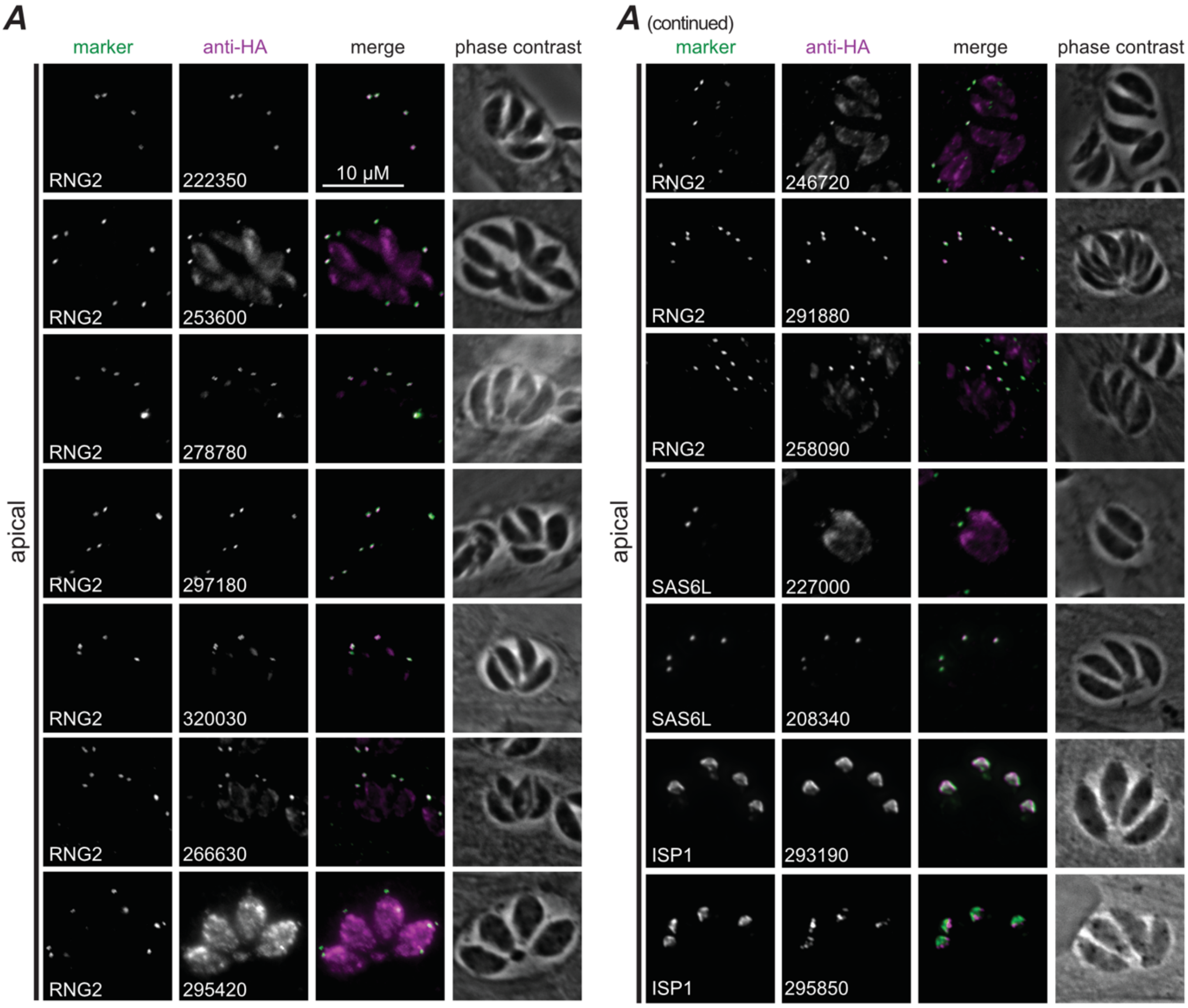
A
A (continued)
marker
anti-HA
merge
phase contrast
apical
10 μM
RNG2
222350
RNG2
253600
RNG2
278780
RNG2
297180
RNG2
320030
RNG2
266630
RNG2
295420
RNG2
246720
RNG2
291880
RNG2
258090
SAS6L
227000
SAS6L
208340
ISP1
293190
ISP1
295850

Figure S2 (continued)

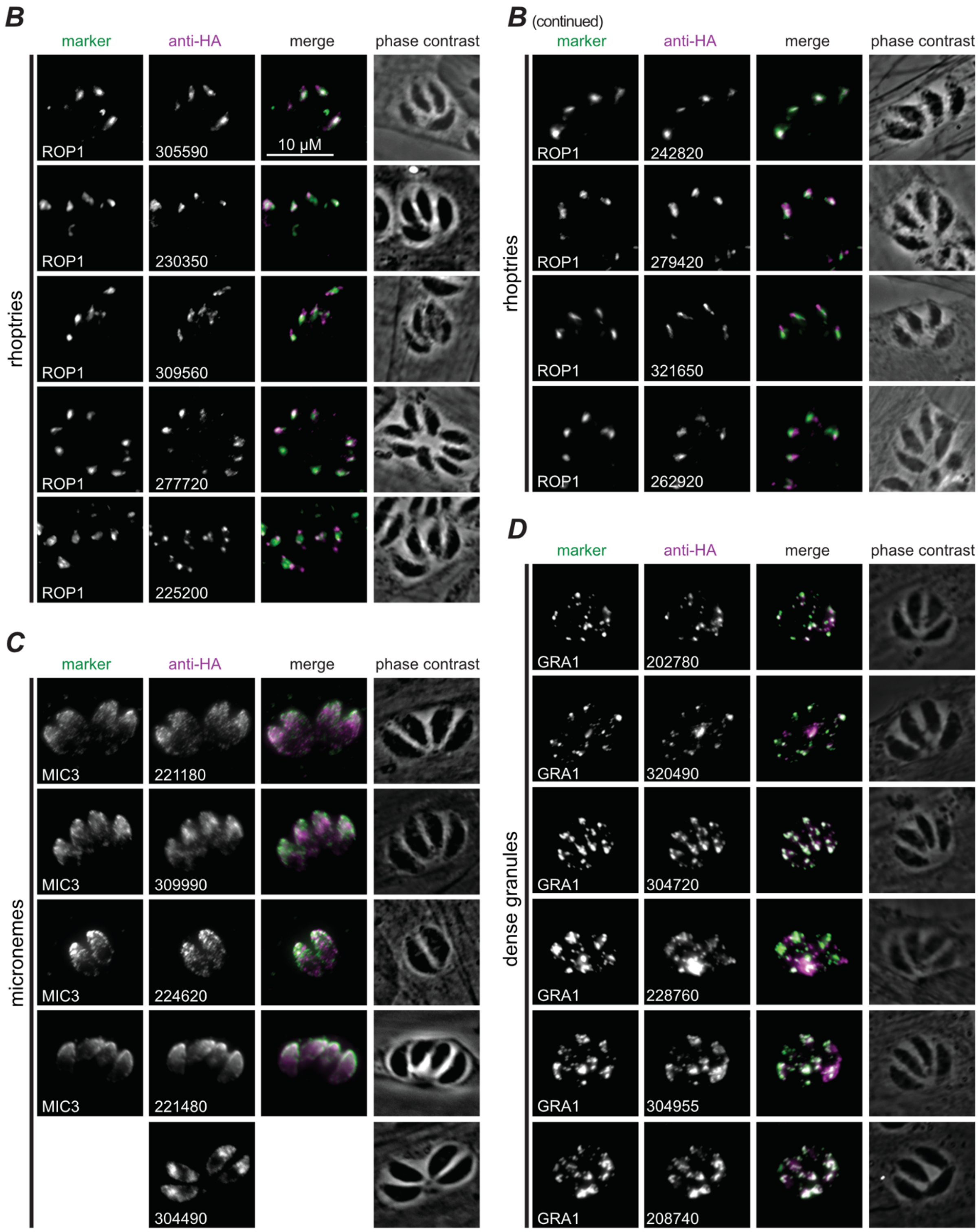

Figure S2 (continued)

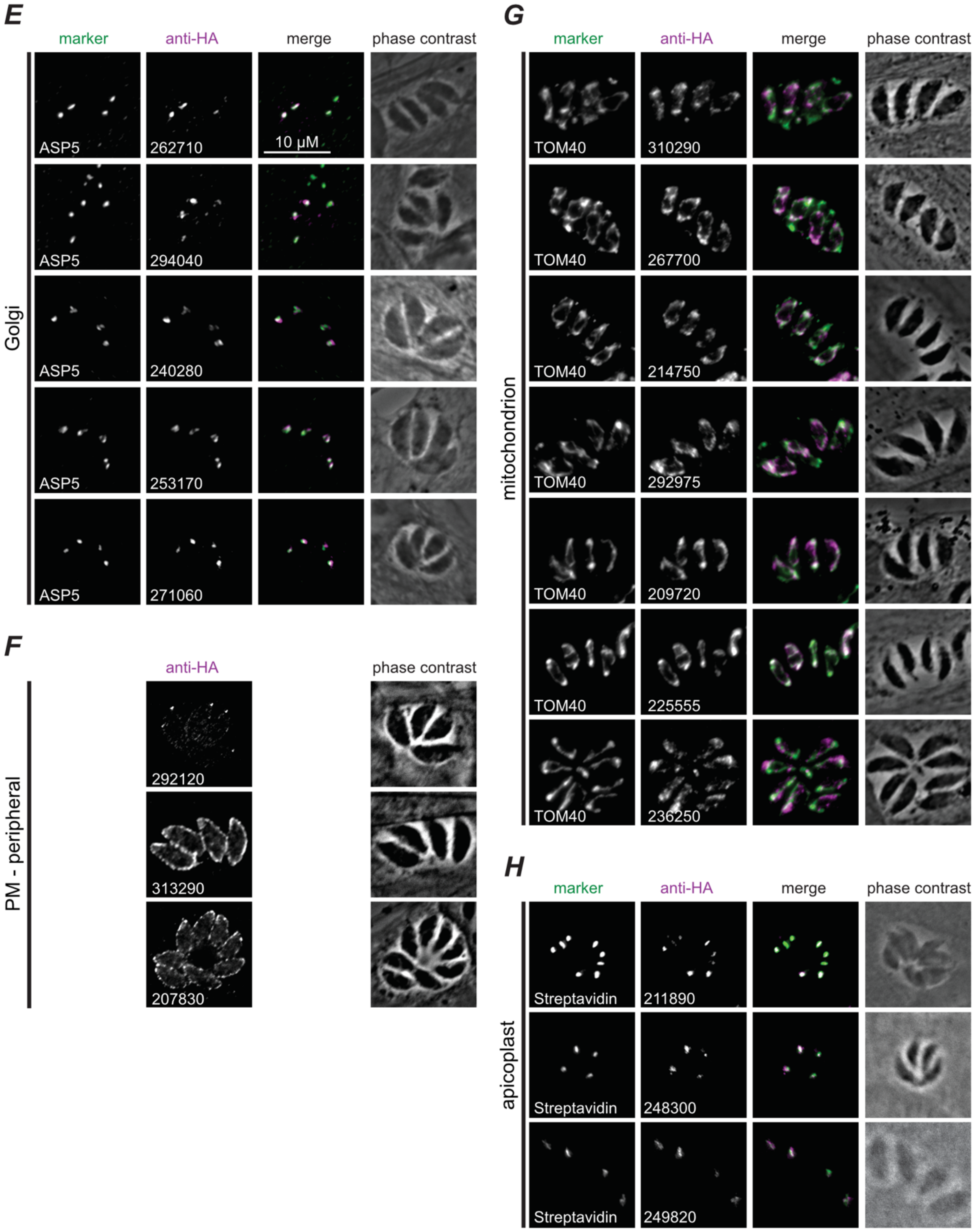


Figure S2. Validation of hyperLOPIT-predicted subcellular locations of select uncharacterized proteins by epitope tagging and immunofluorescence microscopy, continued from Figure 2.

Proteins belonging to the *apical 1* and *2* (***A***), *rhoptries 1* and *2* (***B***), *micronemes* (***C***), *dense granules* (***D***), *Golgi* (***E***), *PM – peripheral 2* (***F***), *mitochondrion-membranes* and *-soluble* (***G***), and *apicoplast* (***H***) clusters are grouped in panels. Scale bar = 10 µm for all.

Figure S3

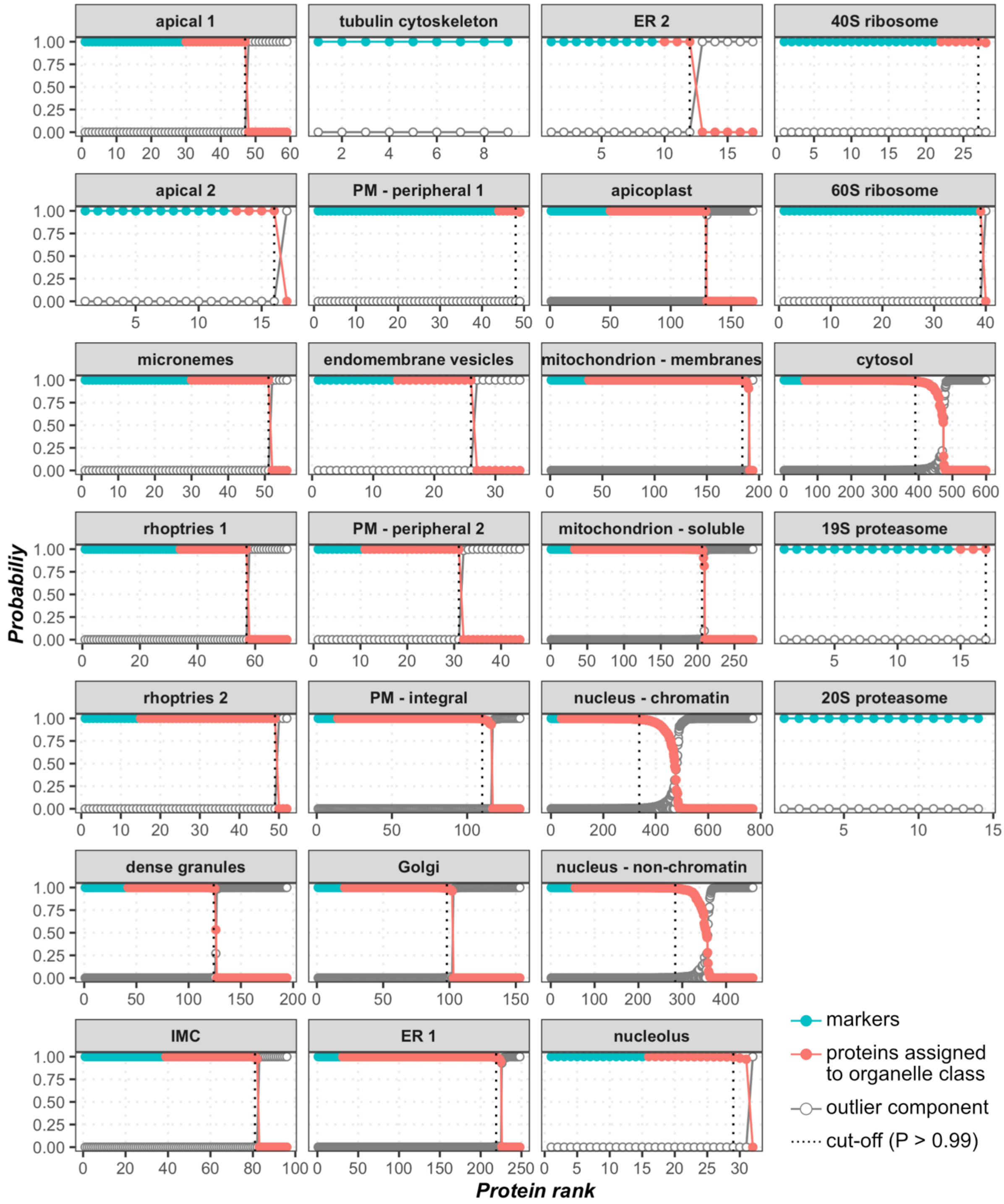


Figure S3. The posterior localization probabilities of 3,832 *T. gondii* proteins determined by a supervised Bayesian classification method TAGM-MAP, related to Figure 3.

Proteins are grouped by the most probable subcellular class, as per the TAGM-MAP classification result, and ranked on the x-axis by their localization probability. The marker proteins are shown in cyan, and the allocated proteins are in red. For each protein, the probability to belong to the outlier component is also shown in grey. The vertical dotted line in each panel indicates the protein localization prediction cutoff (localization probability threshold of 0.99). Only proteins with the localization probability above the threshold of 0.99, i.e. with the rank below the cutoff, retained their class label, whereas the rest of the proteins were labelled as ‘*unassigned*’.

Figure S4

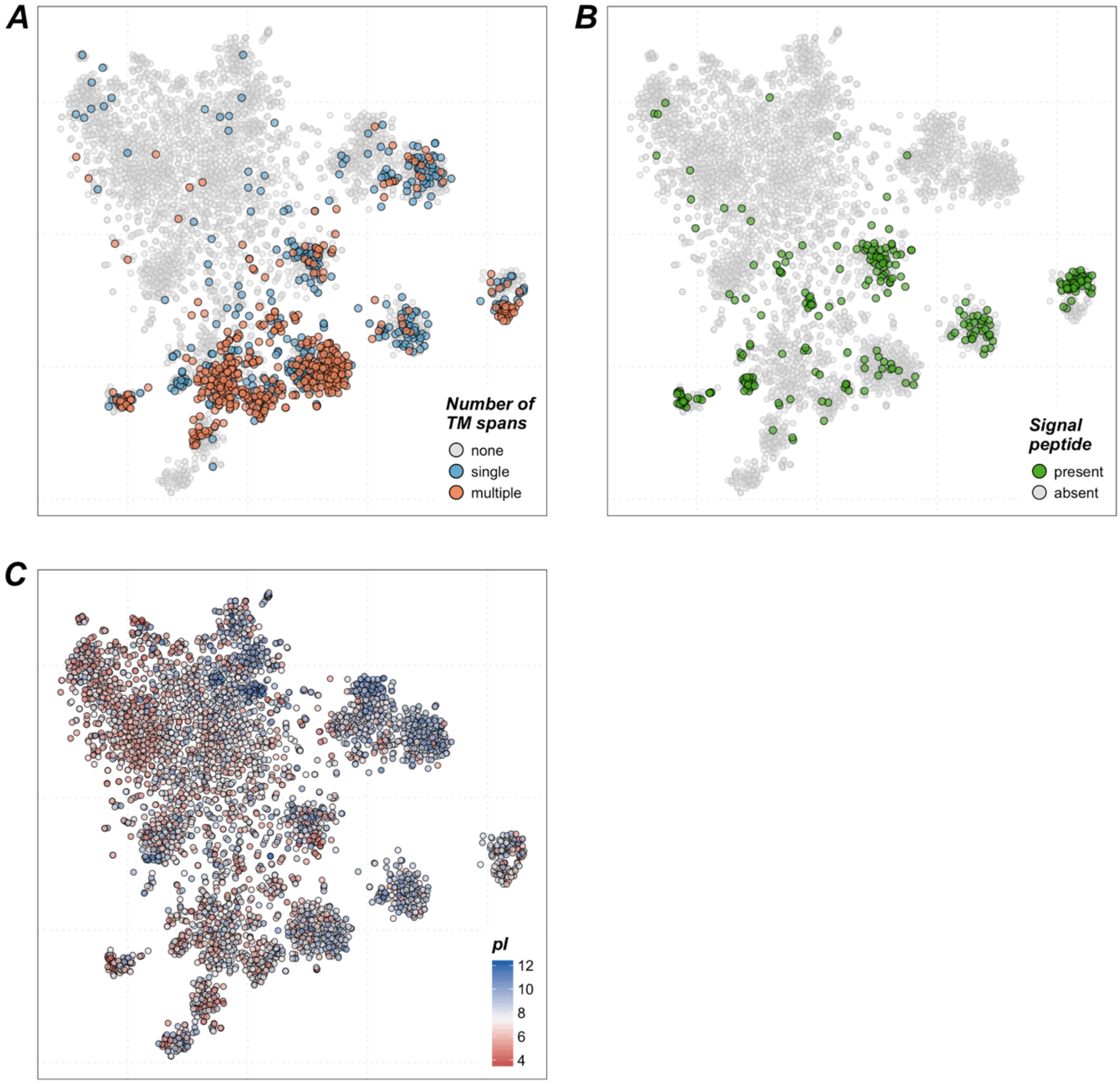


Figure S4. Distributions of select protein sequence features and properties in the spatial proteome of *T. gondii* extracellular tachyzoite, related to Figures 3, and 5.

***A***. A t-SNE projection of the 30plex hyperLOPIT data on 3,832 *T. gondii* proteins with monotopic (blue) and polytopic (red) integral membrane proteins highlighted. TMHMM 2.0 was used to predict transmembrane (TM) spans. The TM spans that overlapped with the signal peptide predicted by SignalP were removed.

***B***. Same as in ***A*** but with the proteins predicted to have a signal peptide (SignalP 5.0) highlighted in green.

***C***. Same as in ***A*** but showing the distribution of protein charge. Proteins are colored according to protein pI computed based on the amino acid sequence. The scale is from red for acidic proteins to blue for basic proteins with the midpoint at pI = 7.4 (colorbar in the bottom-right corner of the panel).

Figure S5

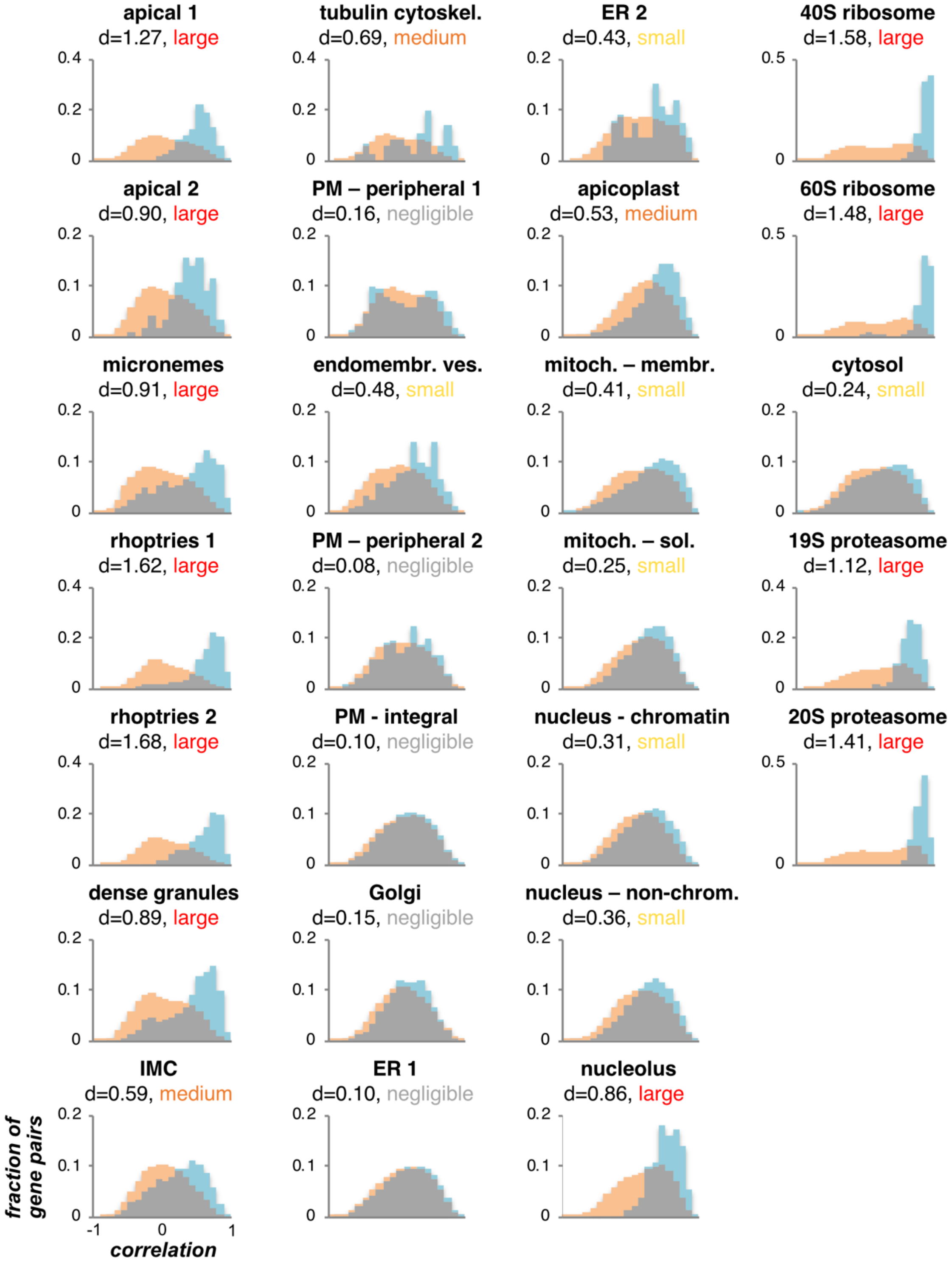


Figure S5. Gene co-expression levels for 26 gene clusters corresponding to *T. gondii* extracellular tachyzoite subcellular compartments determined by hyperLOPIT, related to Figure 4.

The co-expression levels for the genes within hyperLOPIT-defined cluster are shown as light-blue bars in histograms. The co-expression levels between the cluster members and all the genes that are not members of the cluster are shown as orange bars. The Y-axis shows the fraction of gene pairs. The X-axis shows Pearson correlation (the range is from -1 to 1) of non-normalised quantitative transcriptomics data retrieved from ToxoDB.org. Cohen's *d* values with *effect size* descriptors are shown above each plot.

Figure S6

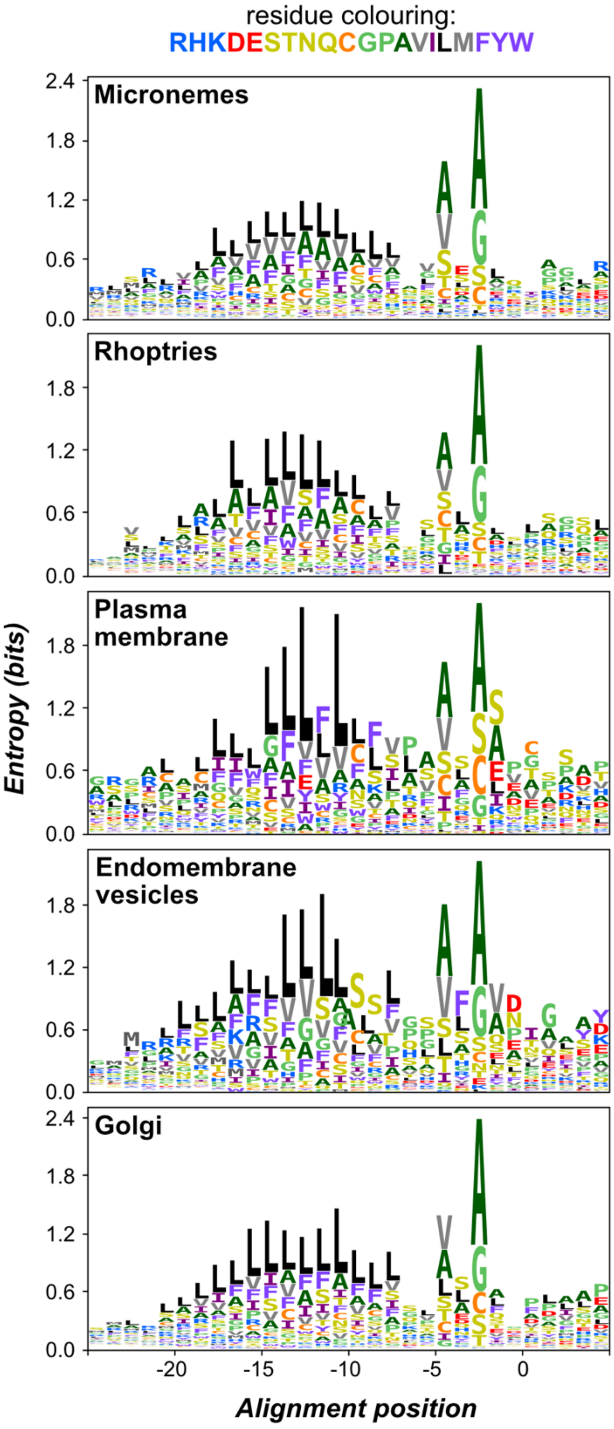


Figure S6. Logo plots of signal peptide (SP) sequences for select cohorts of *T. gondii* proteins, related to Figure 6.

The logo plots show positional abundances of amino acid residue types within and immediately downstream of the SP cleavage site. Proteins from hyperLOPIT-defined *T. gondii* compartment-specific sets and their close homologues from Apicomplexa were aligned anchored at the SP cleavage site (positon 0). Logo plots were generated after randomly sampling 1000 sequences for each data set from position-specific residue abundance probabilities calculated from dissimilarity-weighted sequences. The plots were generated in the same way as in **Figure 5*A***.

Figure S7

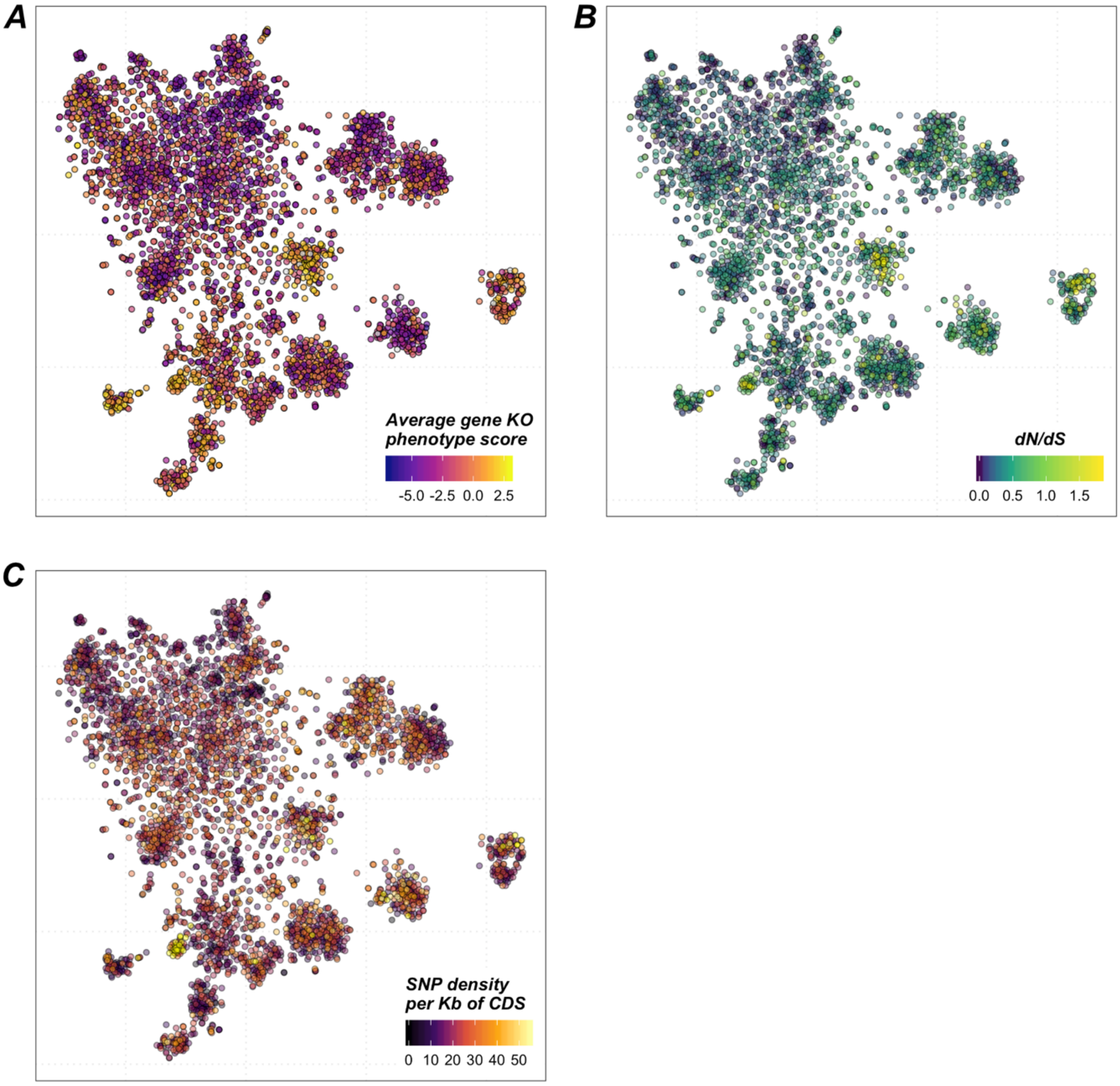


Figure S7. Mapping of the average CRISPR/Cas9-mediated gene knockout phenotype score (A), evolutionary selective pressure (B), and genetic polymorphism (C) on the 30plex hyperLOPIT t-SNE projection of *T. gondii* extracellular tachyzoite spatial proteome data, related to Figure 6.

***A***. Distribution of the average CRISPR/Cas9-mediated gene knockout phenotype score (Sidik et al., 2016). The range is from blue for essential genes to yellow for dispensable genes with the midpoint set at -2.4.

***B***. Distribution of the protein-average ratio of non-synonymous to synonymous point mutations ($d_N/d_S$) (Lorenzi et al., 2016). The scale is clipped at the 99-% quantile of the $d_N/d_S$ range. Data points with extremely high (top-1%) $d_N/d_S$ values are colored in yellow.

***C***. Distribution of the density of single nucleotide polymorphism (SNP) per Kb of protein-coding sequence (CDS) of genes. The SNP density data were retrieved from ToxoDB.org. As in ***A***, the scale is clipped at the 99-% quantile of the data range, with extremely high values (top-1%) shown in yellow.
